# *IRC3* regulates mitochondrial translation in response to metabolic cues in *Saccharomyces cerevisiae*

**DOI:** 10.1101/2021.04.01.438031

**Authors:** Jaswinder Kaur, Kaustuv Datta

## Abstract

Mitochondrial oxidative phosphorylation (OXPHOS) enzymes are made up of dual genetic origin. Mechanism regulating expression of nuclear encoded OXPHOS subunits in response to metabolic cues (glucose vs. glycerol), is significantly understood while regulation of mitochondrially encoded OXPHOS subunits is poorly defined. Here, we show that *IRC3* a DEAD/H box helicase, previously implicated in mitochondrial DNA maintenance, is central to integrating metabolic cues with mitochondrial translation. Irc3 associates with mitochondrial small ribosomal subunit in cells consistent with its role in regulating translation elongation based on Arg8^m^ reporter system. Glucose grown *Δirc3ρ^+^* and *irc3* temperature sensitive cells at 37⁰C have reduced translation rates from majority of mRNAs. In contrast, when galactose was the carbon source, reduction in mitochondrial translation was observed predominantly from Cox1 mRNA in *Δirc3ρ^+^* but no defect was observed in *irc3* temperature sensitive cells, at 37⁰C. In support, of a model whereby *IRC3* responds to metabolic cues, suppressors of Δ*irc3* isolated for restoration of growth on glycerol media restore mitochondrial translation differentially in presence of glucose vs. glycerol.

## Introduction

Mitochondria which are best known as the powerhouse of the cell, requires coordinated gene expression of two spatially distinct genetic material. Mitochondria are essential for an organism’s viability and normal physiology and any disruption in its functioning leads to myriad of cellular defects including cancer (Rutter & Hughes, 2015, Scheper, van der Knaap et al., 2007, Stotland & Gottlieb, 2015). *Saccharomyces cerevisiae* has been an invaluable system for deciphering mitochondrial function, due to its ability to survive without respiration as well as mitochondrial DNA (mtDNA), permitting the characterization of mutants that impair mitochondrial functioning. In *Saccharomyces cerevisiae*, mtDNA encodes for eight proteins, seven of which are transmembrane proteins that are essential components of oxidative phosphorylation (OXPHOS) machinery and one soluble protein Var1, an essential component of mitochondrial small ribosomal subunit. In addition to these eight protein coding genes, mtDNA also encodes for rRNA’s (15S and 21S) and complete set of tRNA’s required for gene expression. Remainder of proteins that make up the OXPHOS subunits, factors required for mitochondrial transcription, translation including mitochondrial ribosomal proteins are encoded by the nuclear genome which are translated in the cytosol and imported into the mitochondria (Fox, 2012).

Translation of mitochondrial mRNA, in addition to general translation factors such as Tuf1 (EF-Tu) and Mef1, Mef2 (EF-G), require membrane bound mRNA specific translation activators (Costanzo, Bonnefoy et al., 2000, Nagata, Tsunetsugu-Yokota et al., 1983, Ott, Amunts et al., 2016, Rasmussen, 1995, Rosenthal & Bodley, 1987, Towpik, 2005, Vambutas, Ackerman et al., 1991). In the absence of either shine-Dalgarno sequences or 5ꞌ cap on mitochondrial transcripts, these mRNA specific translation activators recognize the 5ꞌ UTR of mRNA to localize them to the mitochondrial inner membrane where they aid in loading of membrane bound ribosomes to initiate mitochondrial translation (Brown, Costanzo et al., 1994, Costanzo & Fox, 1993, Dunstan, Green-Willms et al., 1997, Green-Willms, Fox et al., 1998, Haffter, McMullin et al., 1991, Lipinski, Kaniak-Golik et al., 2010, Naithani, Saracco et al., 2003). In fact, each mitochondrial mRNA has their specific set of translation activators (Herrmann, Woellhaf et al., 2013, Kummer & Ban, 2021). Interestingly, altered level or activity of these translation activators are thought to allow yeast cells to sense metabolic cues. Yeast cells when transferred from fermentable to non-fermentable growth leads to enhanced OXPHOS expression and activity. This is brought about by enhanced expression of nuclear encoded OXPHOS subunits and mitochondrial mRNA specific translation factors which upregulates translation of mitochondrially encoded OXPHOS subunits (Couvillion, Soto et al., 2016, Morgenstern, Stiller et al., 2017, Steele, Butler et al., 1996). Another way mitochondrial translation responds to alteration in metabolic cue is by using different mitochondrial translation factors under different conditions. For example, the translation from *COX1* mRNA is controlled by *PET309*, *MSS51* and *MAM33* (Barrientos, Zambrano et al., 2004, Manthey & McEwen, 1995, Perez-Martinez, Broadley et al., 2003, Perez-Martinez, Butler et al., 2009, Roloff & Henry, 2015, Zamudio-Ochoa, Camacho-Villasana et al., 2014). Mam33 is required for the translation of Cox1 at basal level in glucose-grown yeast cells along with Pet309p and Mss51p. However, its function is dispensable in cells adapted for growth in glycerol where *COX1* mRNA translation is regulated primarily by Mss51p and Pet309p (Roloff & Henry, 2015).

In addition to translation factors, optimal mitochondrial gene expression requires the essential function of RNA helicase among other accessory proteins. They regulate every aspect of mitochondrial RNA metabolism including RNA splicing, mRNA turnover/surveillance, translation, and ribosome biogenesis (Bourgeois, Mortreux et al., 2016, Jankowsky & Fairman, 2007, Szczesny, Wojcik et al., 2013). RNA helicases utilize energy released from ATP hydrolysis to promote rearrangements required by nascent RNA molecules to arrive at a mature structure (Chen, Potratz et al., 2008, Henn, Cao et al., 2010, Jarmoskaite & Russell, 2014). These mature structures are achieved by action of helicases to unwind dsRNA, anneal ssRNA, modify RNA-DNA hybrids and displace proteins bound to RNA (Fairman, Maroney et al., 2004, Henn, Bradley et al., 2012, Linder & Jankowsky, 2011). In *Saccharomyces cerevisiae*, the nuclear genome encodes for four mitochondrial helicases which belongs to SFII family of NTP dependent remodelers containing DExH/D motif that regulates different aspects of mitochondrial function. *MSS116* has been shown to play a role in transcription, mRNA splicing and translation (De Silva, Poliquin et al., 2017, Halls, Mohr et al., 2007, Markov, Wojtas et al., 2014, Zingler, Solem et al., 2010), while *MRH4* and *SUV3* are known to be involved in ribosome biogenesis and RNA editing/turnover respectively (De Silva, Fontanesi et al., 2013, Dziembowski, Piwowarski et al., 2003, Malecki, Jedrzejczak et al., 2007, Turk & Caprara, 2010). The fourth mitochondrial helicase, *IRC3* has been implicated in mtDNA recombination and repair (Piljukov, Garber et al., 2020, Sedman, Gaidutsik et al., 2014). However, these studies were carried out in strains deleted for *IRC3*, in which loss of mtDNA could be either due to a direct role of Irc3 in regulating mtDNA recombination/repair or due to an indirect consequence of Irc3’s role in an essential process involving RNA molecules such as transcription and translation.

In the present study, we have shown that *IRC3* is a central regulator of mitochondrial translation. Interestingly, Irc3p regulates mitochondrial translation distinctly in cells grown in carbon source that maintain OXPHOS at basal (glucose) *vs.* elevated levels (galactose/glycerol). Glucose grown *Δirc3ρ^+^* cells were defective for overall mitochondrial protein synthesis in addition to reduction in accumulation of mitochondrial transcript. In contrast, glucose grown cells harbouring *irc3* temperature sensitive mutants have reduced rates of translation, without having any consequence on the levels of mRNA transcript or assembled mitochondrial ribosomal subunit, when shifted to the non-permissive temperature. Interestingly, galactose grown *Δirc3ρ^+^* and cells harbouring *irc3* temperature sensitive mutants, reduction in mitochondrial translation from Cox1 mRNA was observed in *Δirc3ρ^+^* but not in cells harbouring *irc3* temperature sensitive mutants at the non-permissive temperature. Consistent with a role in translation regulation, Irc3 co-fractionates with small ribosomal subunit when cells are grown in either glucose or glycerol. Importantly, we have shown that *IRC3* is required for translation elongation in glucose grown cells. Consistent with a hypothesis whereby mechanism of action and target of *IRC3* are different under conditions that maintain mitochondrial function at a basal level versus conditions that have higher mitochondrial function, we were able to isolate suppressors of *Δirc3ρ^+^* that restored mitochondrial translation to different degrees when grown in glucose versus glycerol.

## Results

### *IRC3* is essential for mitochondrial gene expression

To determine whether *IRC3* regulates an essential mitochondrial function in cells, we sporulated a heterozygous diploid strain of Δ*irc3* generated in the lab. Although, the number of colonies formed on glucose plates in freshly germinated Δ*irc3* or *IRC3* spores were similar, Δ*irc3* spores formed two types of colonies which could be differentiated based on colony size. On glycerol plates fewer number of Δ*irc3* spores formed colonies in comparison to *IRC3* although the colony sizes were comparable. In fact, the number of large colonies formed on glucose plates were equivalent to the number of colonies formed on glycerol plates (Fig. 1A). To minimize the effect of an unlinked mutation on the inability of Δ*irc3* spores to utilize glycerol, Δ*irc3/IRC3* episomally expressing wild type *IRC3* linked to *URA3* were sporulated. The Δ*irc3* spores episomally expressing *IRC3* allele linked to *URA3* were back crossed six times as described in material and methods. The reduced ability of freshly generated Δ*irc3* cells (colonies obtained on 5’FOAD) to utilize glycerol as the sole carbon source was a fraction of the total viable cells in comparison to Δ*irc3* cells ectopically expressing *IRC3* when cultured either in glucose, galactose or glycerol (Fig. S1A). This indicated that ectopic expression of *IRC3* was able to complement the glycerol growth defect. All further experiments were carried out using this strain. Furthermore, the total viable cells that were able to utilize glycerol as the sole carbon source was estimated in Δ*irc3* cells expressing either wild type *IRC3* or vector upon subculturing in glucose for 0hrs and 24hrs. We observed that 10% of progeny cells were competent for cellular respiration in a population of Δ*irc3* cells in comparison to 90% of wild type cells upon sub culturing in glucose or glycerol for 24hrs (Fig. 1B and S1B). Thus, taken together this indicates that *IRC3* is essential for optimal mitochondrial function. Mitochondrial DNA integrity have been reported to be compromised in yeast cells defective for essential processes related to mitochondrial DNA maintenance and gene expression. Measurement of timing of loss of ability to utilize growth on glycerol vs loss of mtDNA in mutant cells is often indicative of the primary role played by the wild type copy of the mutant gene product (Datta, Fuentes et al., 2005, De Silva et al., 2013, Myers, Pape et al., 1985). Thus, we determined the status of mtDNA (*rho* or *ρ*) in a population of haploid Δ*irc3* containing either empty vector or episomally expressing *IRC3* upon subculturing in glucose media for 0hrs and 24 hrs at 30⁰C. Aliquot of cells at each time point were crossed with wild type *ρ⁰* cells of the opposite mating type. Ability of the diploids generated to utilize glycerol are indicative of the presence of mitochondrial DNA in the parental haploid Δ*irc3* cells. Approximately 90-100% of cells episomally expressing wild type copy of *IRC3* retained mitochondrial DNA upon subculturing in glucose. In Δ*irc3* cells, approximately 85% cells retained mtDNA upon subculturing in glucose for 24hrs (Fig 1C). However, further incubation of Δ*irc3* cells led to loss of mitochondrial DNA, such that only 20% of cells retained mitochondrial DNA upon subculturing in glucose for 72hrs (Fig. S1C). Further, to quantify mtDNA and analyse its stability, in *Δirc3* cells expressing either wild type *IRC3* or vector, upon subculturing them in glucose for 0hrs and 24hrs we measured mtDNA copy number with respect to nuclear DNA copy number as described in material and methods. Ratios of mtDNA to nDNA were similar in cells deleted for *IRC3* in comparison to wild type cells even after subculturing in glucose for 24hrs (Fig. S1D). Thus, all subsequent experiments using *Δirc3ρ^+^,* were carried out under culture conditions such that cells retain mitochondrial DNA. Taken together our results indicate that loss of growth on glycerol precedes loss of mitochondrial DNA in Δ*irc3* cells. This is suggestive of defect in an essential process within the mitochondria in Δ*irc3* cells such as those linked to mitochondrial gene expression as the primary cause of loss of cellular respiration while loss of mitochondrial DNA as secondary consequence. Similar phenotypes have been reported for numerous mutant genes that are defective in mitochondrial gene expression (Datta et al., 2005, De Silva et al., 2013, Myers et al., 1985).

**Fig. 1:**
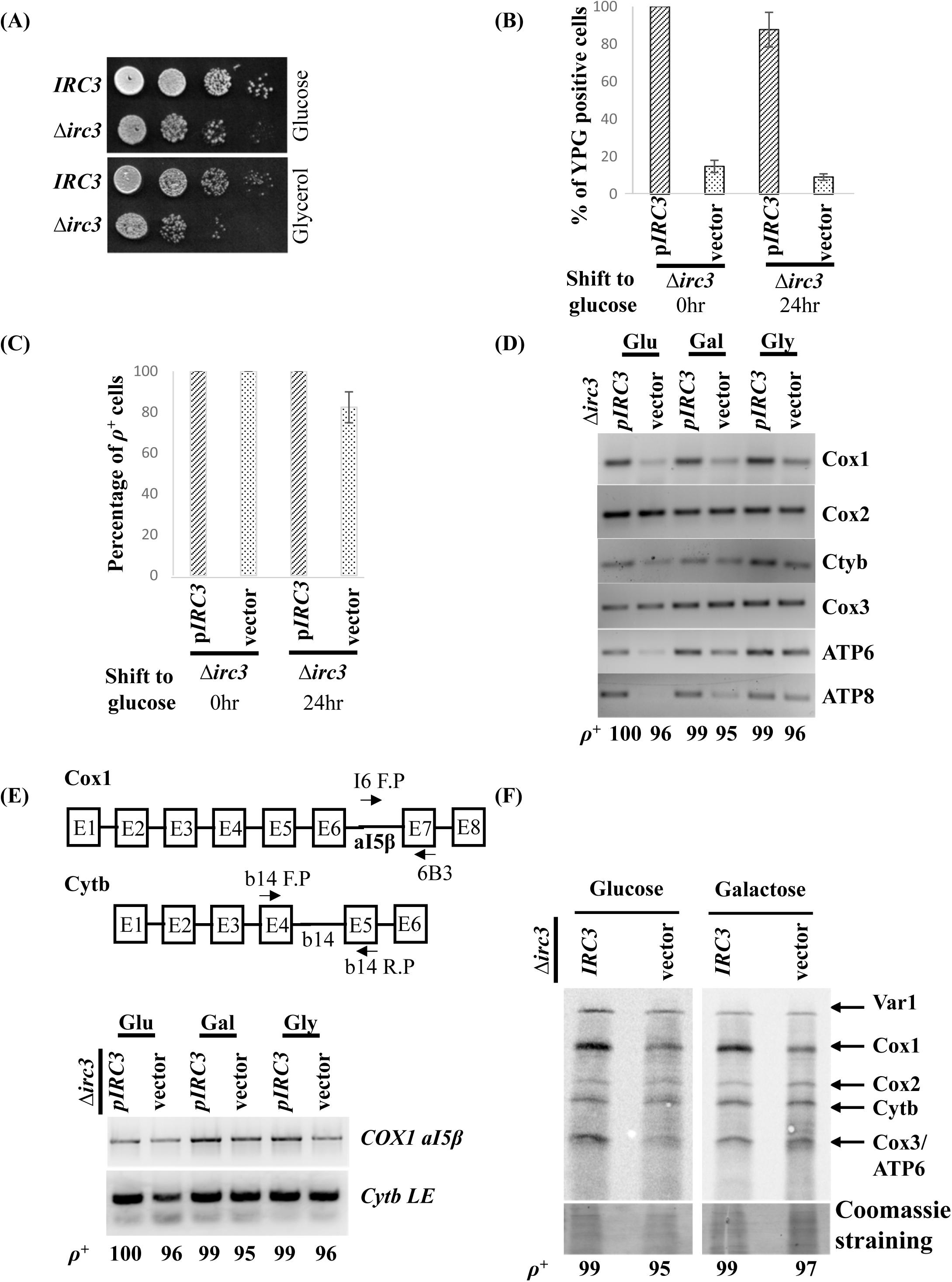
Loss of growth on glycerol precedes loss of mtDNA in Δ*irc3ρ^+^* cells due to aberrant gene expression. (A) Shown are tenfold serial dilution of freshly germinated spores of wild type and Δ*irc3* spotted on glucose or glycerol plates. (B) Percentage of viable progeny cells of Δ*irc3* expressing either episomal copy of *IRC3* or empty vector that can utilize glycerol as the sole carbon source at the indicated time points upon subculturing in glucose. Three independent colonies of wild type and Δ*irc3* were used to calculate the %YPG positive cells. Five independent replicates of this experiment have been performed. (C) Percentage of mtDNA present in progeny of Δ*irc3* expressing either episomal copy of *IRC3* or vector at the indicated time after subculture in glucose. Three independent colonies of wild type and Δ*irc3* were used to calculate the % *ρ^+^* positive cells. Five independent replicates of this experiment have been performed. (D) Transcript levels of mitochondrial encoded genes were assayed in Δ*irc3* cells expressing either episomal copy of *IRC3* or vector grown in glucose, galactose, and glycerol. 21s rRNA and 15s rRNA levels as detected by EtBr staining. Multiple independent biological and technical replicates were carried out and shown are representative images. (E) Shown in the upper panel is a schematic representation of exons and introns in the Cox1 and Cytb transcripts. Labelled by arrows are primers subsequently used to determine the splice variant in Cox1 and Cytb transcripts. Lower panel showed the level of transcript level of pre-mRNA of Cox1(COX1 *aI5β*: product of primer I6 F.P. and 6B3) and Cytb ligated exon (Cytb LE: b14 F.P and b14 R.P). (F) Mitochondrial protein synthesis was measured in Δ*irc3* cells expressing either episomal copy of *IRC3* or vector cultured in either glucose or galactose by incorporation of [^35^S]methionine and cysteine in presence of cycloheximide to inhibit cytosolic protein synthesis. Mitochondria from labelled cells were isolated and equivalent amount of mitochondrial proteins were separated on 17.5% SDS-Page and transferred to nitrocellulose membrane. Radiolabeled proteins were visualized by phosphoimaging. Shown below are coomassie stained gels of radiolabeled mitochondrial protein extracts. Percentage of mitochondrial DNA present in the respective strains are indicated below in (D), (E) and (F).

*IRC3* is a putative helicase of the SFII family, a majority of which have been known to act on RNA molecules (Cordin, Banroques et al., 2006, Fairman-Williams, Guenther et al., 2010). Thus, *IRC3* could potentially be involved in transcription, splicing, turnover, ribosome assembly or translation to regulate mitochondrial gene expression. Mitochondrial transcript levels were examined in Δ*irc3ρ^+^* cells grown either in glucose, galactose, or glycerol by performing reverse transcriptase-PCR as described in material and methods. Transcript levels of *COX1, Cytb*, *ATP6*, *ATP8* and *ATP9* were reduced in Δ*irc3ρ^+^* cells in comparison to wild type cells, although steady state levels of *COX2* and *COX3* mRNA remained unchanged (Fig. 1D). These observations were consistent irrespective of the carbon sources used for cell culture (Fig. 1D). To determine whether the reduced transcript levels in cells deleted for *IRC3* is a consequence of improper intron splicing, we analysed Δ*irc3ρ^+^* cells for efficient splicing of *COX1* and *Cytb* transcripts. To check the accmulation of transcripts where a15β intron is retained in cDNA’s synthesised from *COX1* transcripts primers used for reverse transcriptase-PCR were designed in between a15β and exon7 (I6F.P, 6B3). To amplify the ligated exon from synthesised cDNA for *Cytb* transcripts, primers were designed from exonic regions upstream and downstream of b14 (b14F.P, b14R.P) introns (Fig. 1E top). In comparison to wild type cells, the levels of unspliced mRNA i.e., *COX1 a15β* did not accumulate in Δ*irc3ρ^+^* cells (Fig. 1E). The reduction in levels of *Cytb* ligated exon were similar to reduced transcript levels of *Cytb* (Fig. 1E and 1D). In addition we found that Δ*irc3* cells harboring intronless mtDNA led to similar glycerol growth defects as in Δ*irc3* cells harboing wild type mtDNA (Fig. S1E). These results are suggestive of a role for *IRC3* independent of regulating RNA splicing. To ascertain whether mitochondrial translation is also perturbed in Δ*irc3ρ^+^*, newly synthesized polypeptides were labelled in Δ*irc3ρ^+^* cells either episomally expressing *IRC3* or transformed with vector backbone after growth in glucose and galactose. Growth of cells in glucose lead to glucose repression whereby expression of genes required for alternate carbon sources as well as mitochondrial oxidative phosphorylation are down regulated. However, growth in galactose does not repress genes required for either fermentation or cellular respiration and thus considered to be neutral to these pathways (Conrad, Schothorst et al., 2014, Johnston, 1999, Scheffler, de la Cruz et al., 1998). *De novo* labelling of mitochondrial proteins in Δ*irc3ρ^+^* cells grown on glucose led to an overall reduction in incorporation of [^35^S]methionine and cysteine into newly synthesized proteins in comparison to wild type (Fig. 1F). In contrast *de novo* labelling of mitochondrial proteins in Δ*irc3ρ^+^* cells grown in galactose lead to reduced incorporation of [^35^S]methionine and cysteine into newly synthesized Cox1 polypeptides in comparison to wild type cells (Fig. 1F). This indicates that Irc3p is pivotal for mitochondrial translation irrespective of the type of carbon source the cells are grown in.

Within the mitochondria, transcription and translation are tightly coupled presumably to make the mitochondrial gene expression more efficient and faster (Fontanesi, Soto et al., 2006, Kehrein, Schilling et al., 2015, Rodeheffer, Boone et al., 2001, Singh, Salvatori et al., 2020). Thus, diminished accumulation of mRNA levels in *Δirc3ρ^+^* could be indirect consequences of altered translation rates or *vice versa*.

### *IRC3* is essential for mitochondrial translation during basal OXPHOS activity

To ascertain whether transcription or translation is primarily controlled by *IRC3*, two conditional alleles of *IRC3* designated as *irc3-1* and *irc3-2* were generated. Cells expressing either temperature-sensitive alleles or wild type allele of *IRC3* when cultured in glucose at the permissive temperature of 23⁰C gave rise to progeny that was able to utilize glycerol at the same rate. On the other hand, cells expressing either *irc3-1* or *irc3-2* when sub cultured in glucose at the non-permissive temperature of 37⁰C, gave rise to fewer progenies that can utilize glycerol (Fig. 2A). We confirmed that cells expressing either *irc3-1* or *irc3-2* retained mtDNA copy number similar to wild type levels upon subculturing in glucose for 24hrs at both at 23⁰C and 37⁰C (Figs 2B and S2A).

**Fig. 2:**
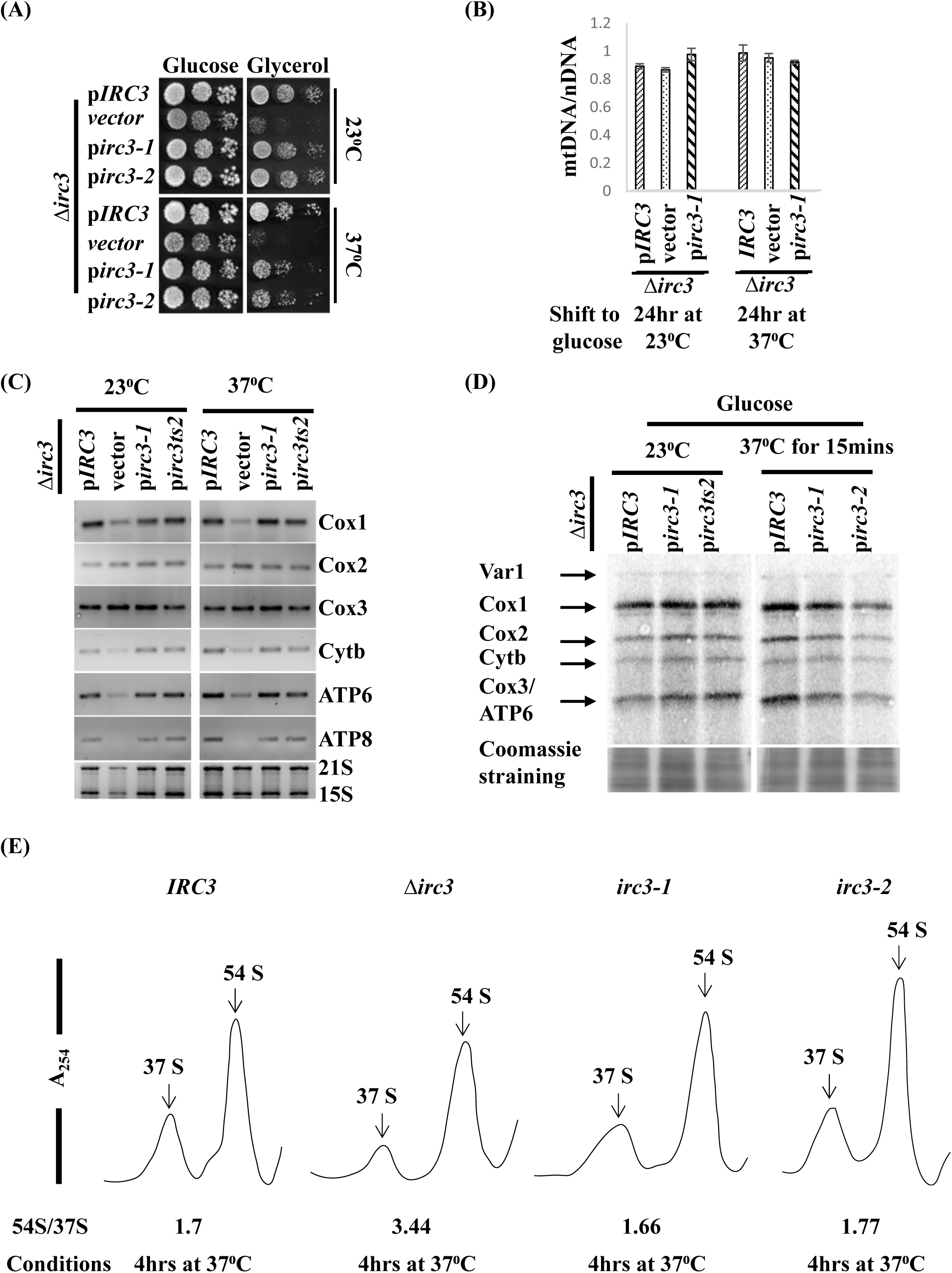
Rapid depletion of functional *IRC3* in cells grown in glucose leads to aberrant mitochondrial translation. (A) Shown are tenfold serial dilution of glucose grown Δ*irc3ρ^+^* cells expressing either episomal copy of *IRC3,* vector *, irc3-1* and *irc3-2* cells, spotted on glucose and glycerol plate and incubated at 23⁰C and 37⁰C. (B) Ratio of nuclear to mitochondrial DNA, was estimated in the indicated strains after sub culturing in glucose for the mentioned time period either at 23⁰C or 37⁰C. (C) Transcript levels of mitochondrial encoded genes were assayed in the indicated strains grown in glucose. 21S rRNA and 15S rRNA levels as detected by EtBr. (D) Mitochondrial protein synthesis was measured by incorporation of [^35^S]methionine and cysteine in the presence of cycloheximide to inhibit cytosolic protein synthesis in the indicated strains grown in glucose at either 23⁰C or after shift to 37⁰C for 15mins. Mitochondria from labelled cells were isolated and equivalent mitochondrial proteins were separated on 17.5% SDS-PAGE. Radiolabeled proteins were visualized by phosphoimaging. Below are coomassie stained gel of radiolabeled mitochondrial protein extracts. (E) Mitochondrial ribosomal subunits from the indicated strains cultured in glucose at 37⁰C for 4hrs were separated on a 10-30% sucrose gradient. RNA content was monitored by taking absorbance at 254nm. The position of 37S and 54S peaks are labeled. The relative ratio of 54S/37S peaks are listed below each profile.

To determine whether *IRC3* primarily regulates transcription, we measured levels of mature transcripts using reverse transcriptase-PCR in cells harbouring either *IRC3* or *irc3-1* or *irc3-2* cultured at either 23⁰C or after a shift to 37⁰C for 2 hrs. We found that levels of mitochondrial mRNA were similar in *irc3-1* and *irc3-2* cells in comparison to wild type cells at 23⁰C or after shifting to 37⁰C (Fig. 2C). These results indicate that reduced steady state levels of mitochondrial mRNA in the strain *Δirc3ρ^+^* in glucose grown cells (Fig. 2C and 1D) could be an indirect consequence of loss of functional Irc3.

To determine whether Irc3p regulates mitochondrial translation, incorporation of [^35^S] labelled methionine and cysteine into newly synthesized mitochondrial proteins were analysed in *IRC3*, irc3*-1*, and *irc3-2* cells grown in glucose, galactose and glycerol. Cells expressing *IRC3, irc3-1* and *irc3-2*, when grown on glucose at 23⁰C showed a similar rate of incorporation of radiolabel into newly synthesized mitochondrial proteins (Fig. 2D). However, when cells were shifted to 37⁰C for 15mins, cells expressing temperature sensitive alleles *irc3-1* and *irc3-2* showed reduced incorporation [^35^S] labelled methionine and cysteine in comparison to wild type cells, indicating the involvement of Irc3 in optimal regulation of mitochondrial translation (Fig. 2D). Interestingly, incorporation of radiolabel in newly synthesized proteins were compromised to a greater extent in *irc3-2* cells (Fig. 2D) which could be due to mutation at a different site than *irc3-1* (Table S3). These results indicate that synthesis of all mitochondrial encoded proteins are affected early upon depleting the functional *IRC3* in cells under growth conditions containing glucose as the carbon source. To rule out the possibility that the protein translation defects observed in *irc3-1* or *irc3-2* mutant cells at 37⁰C were due to an indirect consequence of aberrant mitochondrial ribosomal assembly/stability, the mitochondrial ribosomes isolated from glucose grown cultures of *IRC3, Δirc3ρ^+^, irc3-1* and *irc3-2* cells after shift to 37⁰C for 4 hrs were separated on a sucrose density gradient. The relative ratios of large mitochondrial subunit (54S) to small mitochondrial ribosomal subunit (37S) and their levels were quantified and compared. Levels of mitochondrial small ribosomal subunit (37S) in *Δirc3ρ^+^* cells were severely reduced leading to altered ratios of 54S/37S whereas in wild type, *irc3-1* and *irc3-2* cells similar ratios of 54S/37S subunits and levels were observed upon shift to 37⁰C for 4 hrs (Fig. 2E). This indicated that altered ratios in *Δirc3ρ^+^* are an indirect consequence of disrupted mitochondrial function due to absence of functional Irc3 rather than a direct role for Irc3 during ribosome biogenesis. This also indicates that defective mitochondrial protein synthesis observe in *irc3-1* and *irc3-2* cells at the 37⁰C were not an indirect consequence of defective mitochondrial ribosomal subunit biogenesis.

We have further observed that mitochondrial translation is regulated by *IRC3* differentially depending upon the available carbon source. Equivalent levels of newly synthesized mitochondrial proteins were observed in *IRC3*, *irc3-1* and *irc3-2* cells that had been cultured in glycerol or galactose at 23⁰C or followed by incubation at 37⁰C for 2hrs (Figs S2B and S2C). Consistently, cells expressing *IRC3*, *irc3-1* and *irc3-2* when cultured in glycerol and galactose at 23⁰C or 37⁰C gave rise to progeny that were able to utilize glycerol to the same extent (Figs S2C and S2D). Reduction in levels of *de novo* Cox1p in *Δirc3ρ^+^* (Fig. 1F) is unlikely to be due to defects in ribosome assembly/levels as the ratio of 54S to 37S were similar in *Δirc3ρ^+^* and *IRC3* cells cultured in glycerol for 12hrs at 30⁰C (Fig. S4). Hence, taken together our data supports a role for Irc3 in regulating mitochondrial translation in response to metabolic cues.

### Irc3p fractionates with small mitochondrial ribosomal subunit independent of metabolic cues

Given that *IRC3* is a predicted RNA helicase and reduction in functional *IRC3* in cells leads to aberrant mitochondrial protein synthesis, suggests a role for Irc3p in mitochondrial translation initiation and elongation, either of which would require Irc3p to be associated with the mitochondrial ribosome. In absence of antibodies to the endogenous protein, to detect Irc3 within the mitochondria, we tagged *IRC3* with 13Xmyc at the C-terminus. We confirmed that Irc3-myc encodes for a functional protein localized to the mitochondria (Fig. S3A and S3B). Interestingly, we had observed that mitochondrial translation was perturbed to different levels in cells expressing mutant *IRC3* when cultured under conditions that maintain mitochondrial OXPHOS function at basal or elevated level (Fig. 1F, 2D, S2B and S2C). Thus, we determined whether Irc3-myc physically associates with mitochondrial ribosomes in exponentially growing cells cultured in glucose and glycerol. In addition, we examined the association of Irc3-myc with mitochondrial ribosomes in cells that had attained stationary phase. Mitochondrial ribosomes were separated on a 10-30% sucrose gradient as described in material and methods and individual fractions were probed for the presence of Mrp7 and Mrp13 to ascertain the fractions containing 37S small subunit and 54S large subunit. Irc3-myc peaked in fractions containing Mrp13 in mitochondria from exponentially growing cells cultured in glucose and glycerol (Fig. 3A). A small proportion of the total signal from Irc3-myc also co-fractionated with both Mrp7 and Mrp13 indicating an association with 74S in mitochondria from exponentially growing cells cultured in glucose and glycerol (Fig. 3A). In contrast when mitochondrial ribosomes were separated on a sucrose gradient from stationary phase cell cultured in glycerol, majority of Irc3-myc was found at the top of the gradient and only a small fraction of Irc3-myc pool were present in fractions containing Mrp7, followed by minor pool in heavier fractions (Fig. 3B).

**Fig. 3:**
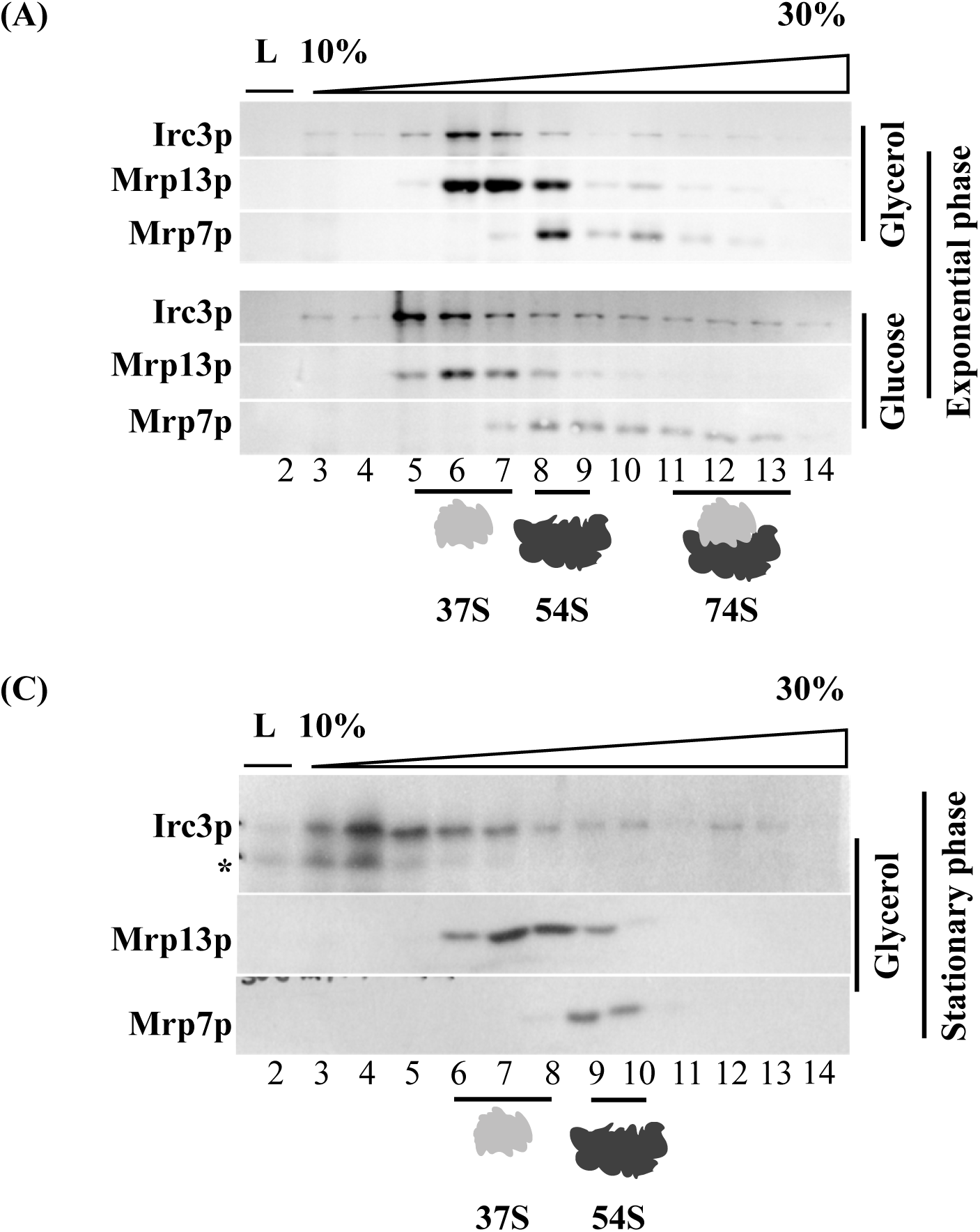
Irc3p associates with small subunit of mitochondrial ribosome during exponential growth phase. Mitochondrial ribosomes from cells expressing functional *IRC3-Myc*, which are cultured either in (A) glucose or glycerol to exponential phase or (B) in glycerol to stationary phase were separated by ultra-centrifugation on a 10-30% sucrose gradient containing 400mM NH_4_Cl at 135,000g for 4hrs. Equivalent protein fractions were TCA precipitated and separated on SDS-PAGE and subject to immunoblot analysis using antibodies against c-Myc, Mrp7p and Mrp13p. The position of 37S, 54S and 74S are shown at the bottom. (*) This is a cross reacting signal obtained from anti-Myc antibody likely to be a cleavage product of Irc3 during sample processing.

### Irc3p is involved in mitochondrial translation elongation

Absence or reduction in functional *IRC3* led to reduced mitochondrial protein synthesis. This could be due to a defect in either translation initiation or elongation or both. To directly examine role of *IRC3* in translation initiation from mitochondrial mRNA, we used engineered strains YC162, RG140 and RG139, carrying a mitochondrial reporter gene in Δ*irc3* expressing either *IRC3* or vector or *irc3-1* or *irc3-2*. In these strains Arginine8 (Arg8), an essential matrix localized enzyme required for arginine biosynthesis has been deleted from the nuclear genome and a recoded version termed Arg8^m^ is expressed from the mitochondrial DNA, specifically replacing the coding sequence of Cox1, Cox2 and Cox3 keeping the 5’ and 3’ UTRs intact respectively (Fig. S5)(Mays, Camacho-Villasana et al., 2019). Functional mitochondrial translation initiation in these strains can be scored based on the ability of cells to grow on synthetic plates lacking arginine (SD-Arg). YC162, RG140 and RG139 deleted for *IRC3* showed growth defect on SD-Arg consistent with reduced Arg8^m^ protein accumulation in mitochondria (Fig. 4A and B). However, engineered Δ*irc3* cells episomally expressing either *irc3-1* and *irc3-2* allele showed similar growth rates to wild type cells on SD-Arg at both permissive and non-permissive temperature (Fig. 4A). Consistently, steady state levels of Arg8^m^ in mitochondria from cells cultured at non-permissive temperature were also similar in these cells (Fig. 4B). These results indicate that although *IRC3* is a key regulator of mitochondrial translation, defects observed after rapid depletion of functional *IRC3* does not severely compromise the initiation process.

**Fig. 4:**
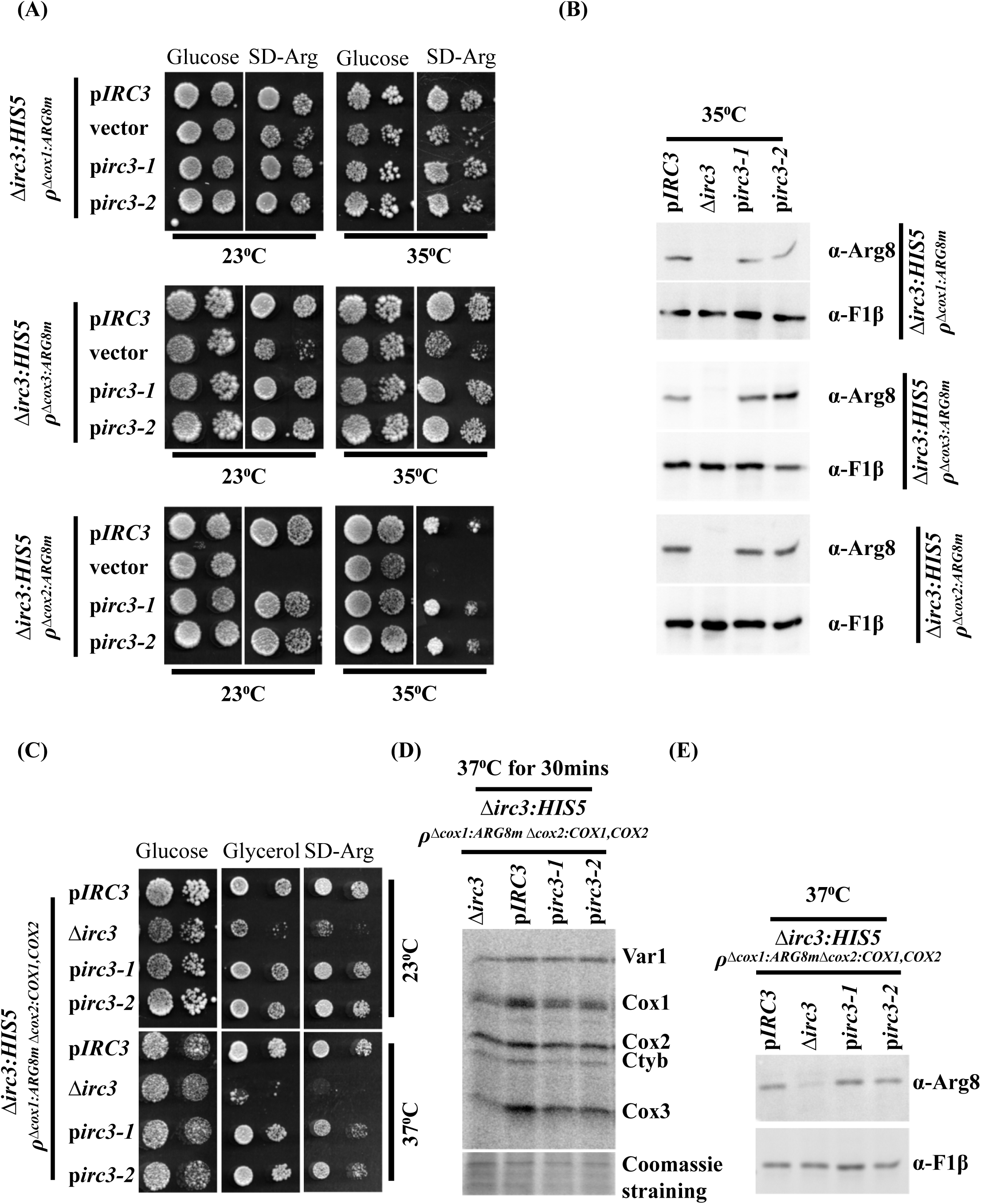
Irc3p regulates mitochondrial translation elongation and not initiation. (A) Shown are serial dilution ofΔ*irc3ρ^+^*cells expressing either wild copy of *IRC3*, vector, *irc3-1* or *irc3-2* harboring mtDNA modified to carry Arg8^m^ reporter cassettes in place of *COX1, COX2* and *COX3* open reading frames were cultured in glucose prior to spotting on glucose and synthetic media lacking arginine and incubated at the indicated temperature. (B) Steady state levels of mitochondrial Arg8^m^ were measured by immunoblot analysis in the indicated strains at non-permissive temperature. Anti-F1β was used as a loading control. (C) Shown are serial dilution of Δ*irc3ρ^+^*cells expressing either wild copy of *IRC3*, vector, *irc3-1* or *irc3-2* harboring mtDNA modified to carry Arg8^m^ reporter cassettes in place of COX1 ORF, COX1 and COX2 ORF were placed under COX2 5’ and 3’ UTR cultured in glucose prior to spotting on glucose, glycerol and synthetic media lacking arginine and incubated at the indicated temperature. (D) Mitochondrial protein synthesis was measured in the indicated strains upon shift from 23⁰C to 37⁰C for 30mins by incorporation of [^35^S]methionine and cysteine in presence of cycloheximide to inhibit cytosolic protein synthesis. Mitochondria from labelled cells were isolated and equivalent amount of proteins were separated on 17.5% SDS-PAGE. Radiolabeled proteins were visualized by phosphoimaging. Below are coomassie stained gel of radiolabeled mitochondrial protein extracts. (E) Steady state levels of mitochondrial Arg8^m^ and F1β were measured in the indicated strains cultured at 37⁰C.

To differentiate the role of *IRC3* more specifically in mitochondrial translation initiation and elongation, we used the engineered strain XPM171a and XPM78a a mitochondrial reporter gene in Δ*irc3ρ^+^* expressing either *IRC3* or vector or *irc3-1* or *irc3-2*. In XPM171a, recoded Arg8^m^ gene was placed under the control of *COX1* 5’ and 3’UTR while *COX1* and *COX2* ORFs were placed under the control of *COX2* 5’ and 3’UTR (Fig. S5) (Perez-Martinez et al., 2003). This allows translation initiation to be monitored as a function of growth on SD-Arg and translation elongation to be scored as a function of growth on glycerol (YPG). When cells were cultured in glucose medium, Δ*irc3* cells in comparison to wild type cells were severely defective for growth on SD-Arg and YPG. (Fig. 4C). In contrast, Δ*irc3* cells episomally expressing either *irc3-1* or *irc3-2* allele were defective for growth on YPG but not SD-Arg at 37⁰C in comparison to *IRC3* cells while at 23⁰C *IRC3*, *irc3-1* and *irc3-2* grew at similar rates on both SD-Arg and YPG (Fig. 4C). Consistent with a defect in translation elongation in glucose cultured cells we found that incorporation of [^35^S] labelled methionine and cysteine into newly synthesized Cox1 as well as other mitochondrial encoded proteins were reduced in Δ*irc3ρ^+^*, *irc3-1* and *irc3-2* cells upon shift to 37⁰C from 23⁰C for 30 mins (Fig. 4D). Consistent with no significant defects in translation initiation Δ*irc3ρ^+^* cells episomally expressing either *IRC3* or *irc3-1* or *irc3-2* at 37⁰C accumulated similar levels of Arg8^m^ in the mitochondria (Fig. 4E). XPM78a expresses 512 nucleotides of intron less *COX1* fused to Arg8^m^ under the control of *COX1* 5’ and 3’UTR (Fig. S5A). Within the Arg8^m^ sequence, present is the cleavage site for pre-Arg8 such that Cox1 and Arg8^m^ function independently (Perez-Martinez et al., 2003). This allows translation elongation to be scored on SD-Arg and glycerol plates. Cells harbouring Δ*irc3* allele was compromised for growth on both SD-Arg and glycerol in comparison to wild type (Fig. S5B). Consistently incorporation of [^35^S] labelled methionine and cysteine into newly synthesized Cox1 and Arg8^m^ in Δ*irc3* cells were reduced in comparison to wild type cells(Fig. S5B). Growth defect in Δ*irc3ρ^+^*cells expressing *IRC3, irc3-1* and *irc3-2* on SD-Arg and glycerol plates could not be ascertained as there was a drastic loss of cell viability at the non-permissive temperature of 37⁰C when incubated for greater than six hours (data not shown). In addition, we were not able to visualise Cox1-Arg8 fusion product, likely due to its labile nature. Taken together these results indicate that Irc3p is involved in mitochondrial translation elongation.

### Rho suppressors of *Δirc3,* restores translation differentially based on metabolic cues

We have shown that Irc3p regulates mitochondrial translation differently in cells based upon the available carbon source for growth. To study the molecular mechanism by which Irc3 regulates mitochondrial function, we isolated spontaneous suppressor of Δ*irc3ρ^+^* using two independent strategies. In the first screen, suppressor of Δ*irc3 ρ^+^* were identified by plating Δ*irc3ρ^+^* for single cell colonies on solid media containing glycerol. Cells that were able to utilize glycerol at a faster rate were initially scored as suppressors. Two suppressors of Δ*irc3* (*Δirc3ρ^SUP1^* and *Δirc3ρ^SUP2^)* that were able to maintain growth in respiratory media after repeated subculturing in glucose were analysed further (Fig. 5A). A second independent screen used *Δirc3ρ^+^* strain carrying an unlinked *ade2* mutation in the background. Strains carrying an *ade2* mutation are red in colour when grown in glucose as long as their mitochondrial function are intact as are *IRC3,ade2* (Fig. 5A). Strains with compromised mitochondrial function that are deficient for respiration are white in colour ((Kim, Sikder et al., 2002, Sesaki & Jensen, 2001)) as are i.e., *Δirc3,ade2* (Fig. 5A). To find second site suppressors of *Δirc3,ade2* cells were plated at on solid glucose media and scored for red colonies sectors. Four suppressors (*Δirc3,SUP3ρ^+^, Δirc3ρ^SUP4^, Δirc3ρ^SUP5^,* and *Δirc3ρ^SUP6^)* were identified and further confirmed for growth on glycerol media (Fig. 5A). Interestingly, the suppressors identified by the first screen formed white coloured colonies on glucose plates in spite of the presence of an *ade2* mutation and on glycerol plates colony size was larger than wild type (Fig. 5A).

**Fig. 5:**
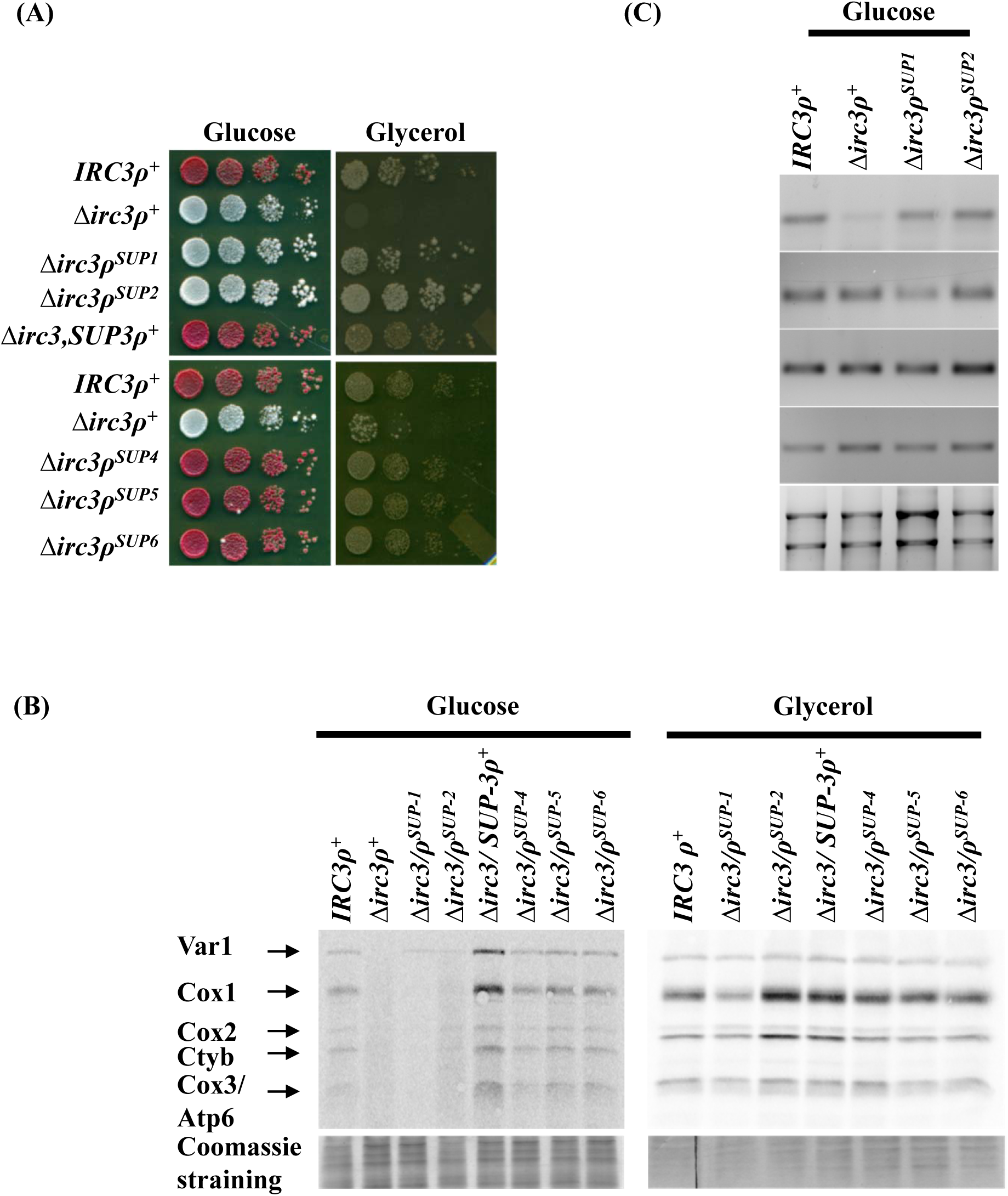
Restoration of mitochondrial protein synthesis in Δ*irc3ρ^+^* suppressors are dependent on carbon source used for cell growth. (A) Shown are tenfold serial dilution of wild type, Δ*irc3ρ^+^*and Δ*irc3ρ^+^* suppressors spotted on glucose and glycerol plates. (B) Mitochondrial protein synthesis was measured in wild type, Δ*irc3ρ^+^* and the indicated suppressors of Δ*irc3ρ^+^* cultured in glucose and glycerol respectively by incorporation of [^35^S]methionine and cysteine in presence of cycloheximide to inhibit cytosolic protein synthesis. Mitochondria from labelled cells were isolated and proteins were separated on 17.5% SDS-PAGE. Radiolabeled proteins were visualized by autoradiography. Below are Coomassie stained gel of radiolabeled mitochondrial extracts. (C) Transcript levels of the indicated mitochondrial encoded genes were assayed in wild type, Δ*irc3ρ^+^, Δirc3ρ^SUP1^*and Δ*irc3ρ^SUP2^* under glucose culture conditions. 21s rRNA and 15s rRNA levels are detected by EtBr staining.

Diploids generated by crossing the suppressor strains with *Δirc3ρ^0^* retained the ability to grow on glycerol, thereby indicating that the suppressor mutation was dominant in character. Among the six suppressors identified five suppressors contained mutation in the mitochondrial genome and one suppressor contained mutation in the nuclear genome as deciphered based on the crosses described in (Fig. S6). As we have shown before Irc3 regulates mitochondrial translation differentially in glucose grown cultures versus galactose/glycerol grown cultures, we determined whether mitochondrial translation was restored to the same levels in all suppressors cultured in glucose or glycerol. We found that incorporation of [^35^S] labelled methionine and cysteine into newly synthesized mitochondrial proteins were restored to wild type levels in *Δirc3,SUP3ρ^+^, Δirc3ρ^SUP4^, Δirc3ρ^SUP5^*and *Δirc3ρ^SUP6^* independent of carbon source in the growth medium (Fig 5B). Interestingly in *Δirc3ρ^SUP1^*and *Δirc3ρ^SUP2^* mitochondrial translation was restored to nearly wild type levels when glycerol was used as a carbon source in growth media. However mitochondrial translation was not restored to wild type levels when glucose was used as a carbon source in growth media (Fig 5B). Levels of mitochondrial mRNA were similar in *Δirc3ρ^SUP1^* and *Δirc3ρ^SUP2^* suppressors in comparison to wild type cells grown in glucose indicating that reduced translation levels in glucose grown cells were not due to reduced transcript levels of mitochondrial mRNA (Fig 5C). These results further indicate that Irc3 regulates mitochondrial translation, and the mode of regulation is dependent on the carbon sources available for cell growth.

## Discussion

Translation is a complicated process involving steps of translation initiation and peptide elongation followed by termination and ribosome recycling. Translation rates are regulated by numerous protein factors and mRNA secondary structures that vary with environmental conditions (Dever, Kinzy et al., 2016). In mitochondria, the translation system has an added layer of complexity where the machinery is tightly associated by the mitochondrial inner membrane and requires membrane embedded mRNA specific translation activator for protein synthesis (Fox, 2012). These presumably are the reason why a membrane free *in vitro* translation system has not yet been developed and the mechanism underlying mitochondrial protein synthesis remains poorly understood.

In this study we used genetic and biochemical methods to investigate the role of putative DExH/D helicase *IRC3,* which could be speculated to modify structures involving either DNA-DNA or RNA-RNA or DNA-RNA hybrids during mitochondrial DNA synthesis/repair and gene expression respectively. We observed that cells deleted for *IRC3* rapidly lose the ability to utilize glycerol as the sole carbon source followed by relatively slow rate of mitochondrial DNA loss (Fig.1), consistent with previously reported examples of proteins required for mitochondrial gene expression (Datta et al., 2005, De Silva et al., 2013, Myers et al., 1985). Furthermore, we demonstrate as expected for a protein to regulate mitochondrial translation Irc3-myc associate with mitochondrial small ribosomal subunits (Fig. 3A) and deletion of *IRC3* led to decreased mitochondrial translation (Fig.1). Although the precise molecular details by which Irc3 regulates mitochondrial translation is unknow, our results support a model where Irc3 regulates translation elongation. First, upon direct examination of mitochondrial translation in *irc3-1* and *irc3-2* cells we observed a decrease in *de novo* polypeptide synthesis upon shift to the non-permissive conditions within the first 15 mins (Fig. 2). Secondly, using Arg8^m^ reporter gene, we observed that Irc3 is required specifically for translation elongation (Fig. 4 and Fig S5B).

It is well known that in response to metabolic cues (glucose versus galactose/glycerol) yeast cells differentially regulate gene expression (Brauer, Huttenhower et al., 2008, DeRisi, Iyer et al., 1997, Gasch, Spellman et al., 2000, Paulo, O’Connell et al., 2015, Turcotte, Liang et al., 2010). Culturing cells either in glucose or galactose/glycerol leads to differential utilization of mitochondrial OXPHOS for ATP production. In glucose, mitochondrial OXPHOS activity is maintained at the basal levels while in galactose/glycerol this activity is elevated (Bouchez, Hammad et al., 2020, Takeda, 1981). Shift of yeast cells from glucose to galactose/glycerol leads to increased translation of nuclear and mitochondrial genome encoded OXPHOS subunits while enhanced mitochondrial protein translation is brought about by enhanced expression of mRNA specific translation activators (Couvillion et al., 2016, Morgenstern et al., 2017, Ohlmeier, Kastaniotis et al., 2004). Could additional molecular factors with novel mechanism exist that modulate mitochondrial translation in response to carbon source? We found that Irc3 differentially regulates mitochondrial translation when cells are cultured under conditions which require OXPHOS at a basal versus elevated levels (Figs 1,2 and S2). When glucose is used as the carbon source, protein translation in *Δirc3ρ^+^* mitochondria is reduced from all transcripts whereas when galactose is used as the carbon source protein translation from Cox1 mRNA is predominantly reduced. This contrasts with the observation that accumulation of mitochondrial transcript was equivalently disrupted irrespective of carbon source used for growth (Fig 1). When *irc3-1* and *irc3-2* cells are cultured in glucose at the restrictive temperature rates of protein translation mirrored protein translation rates from *Δirc3ρ^+^* cells. However, when *irc3-1* and *irc3-2* cells were cultured in galactose/glycerol at the restrictive temperature, mitochondrial translation was not altered, and neither were their progeny cells defective for growth on glycerol. Coincidentally, our screening for temperature sensitive mutants was biased towards disrupting *IRC3* function specifically in glucose cultured cells. Thus, taken together one can speculate that Irc3p differentially regulates mitochondrial translation in response to metabolic cues.

Intriguingly, suppressors of *Δirc3ρ^+^* that were isolated restored mitochondrial translation to different extent based on carbon source added in growth medium. Suppressors of *Δirc3ρ^+^* that were isolated for restoration of mitochondrial function on glycerol, restored mitochondrial translation to wild type levels only when glycerol and not glucose was the carbon source added in growth medium. In contrast suppressors of *Δirc3ρ^+^* that were isolated for restoration of mitochondrial function on glucose, restored mitochondrial translation to the same extent irrespective of whether glucose or glycerol were provided for cell growth (Fig. 5). Although it is not possible to comment on the exact nature of the suppressor mutation(s), however they lead to increased processivity of the translation machinery. Mutation(s) on mtDNA in glucose derived suppressors likely increases the translation processivity to a greater extent than glycerol derived suppressors. Could this difference in translation processivity in suppressors of *Δirc3ρ^+^* hypothetically indicate existence of altered mRNA structures of varying degrees of compactness in mitochondrial transcript in cells grown in glucose versus glycerol? It is not completely understood how mRNA’s fold into compact structures in the cell. However, secondary, and tertiary structures in the 5’UTR, coding sequence (CDS) and 3’UTR are well described regulators of essential process such as translation initiation, translation elongation, ribosomal traffic jams and mRNA stability (reviewed extensively in (Jacobs, Mills et al., 2012, Khong & Parker, 2020, Mauger, Siegfried et al., 2013, Mortimer, Kidwell et al., 2014)). Ribosome processivity during the process of translation elongation is believed to be the principal mechanisms that leads to its secondary structure resolution within the CDS (Adivarahan, Livingston et al., 2018, Khong, Matheny et al., 2017, Yu, Meng et al., 2019). Interestingly in glucose grown yeast cells mitochondrial transcript are abundant although their translation is at the basal level. Upon shift to glycerol increase in translation rates are greater than mRNA synthesis rates (Amiott & Jaehning, 2006, Couvillion et al., 2016). Thus, one could imagine that compact mRNA molecules contribute to the previously described basal translation rates in glucose. Conversely, when glycerol is the carbon source for the cell growth, less compact mitochondrial mRNA structures exist, possibly because of enhanced level of association with the ribosomes, requiring Irc3 for translation of a subset of mRNA such as for Cox1.

Thus, how could Irc3p differentially regulate mitochondrial translation when associated with mitochondrial small ribosomal subunit in both glucose and glycerol? Two non-exclusive hypothetical models could be invoked to address this question both of which require Irc3 associating with different interacting partners. These interacting partners could be proteins whose expression is controlled by metabolic cues such as those that have been described as a part of large-scale proteome remodelling that occur when cells are shifted from glucose to galactose/glycerol. In the first model these protein partners in association with Irc3 target the regions of mRNA with compact local secondary RNA structures that are maintained in the mitochondria as a function of their growth condition which require unwinding during translation elongation. An alternate model could be that these distinct interacting partners for Irc3 are associated with the compact mRNA structures. During the process of translation elongation Irc3 bound to the small subunit is responsible for displacing these proteins thereby melting the structures required for translation elongation. Future experiments targeted to parsing the mechanism by which Irc3 regulates mitochondrial translation will require these hypothetical models to be accounted for.

Deletion of *IRC3* has previously been reported to accumulate double stranded mitochondrial DNA breaks (Sedman et al., 2014). ATPase activity of Irc3 protein is essential *in vivo* and *in vitro* is stimulated by DNA molecules especially those mimicking recombination intermediates (Gaidutsik, Sedman et al., 2016, Piljukov et al., 2020, Sedman et al., 2014). Here in we have clearly shown that *IRC3* is required for mitochondrial translation *in vivo*. It is not well understood why disruption of mitochondrial translation often leads to loss of mitochondrial DNA. Is it possible that SFII DExH helicases can target both mtDNA as well as mitochondrial mRNA thereby forming a link between optimal translation and mtDNA stability? Among the four annotated SFII DExH helicase, *SUV3*, which is the RNA helicase component of mitochondrial exoribonuclease complex in addition to its role in RNA turnover (Dziembowski et al., 2003, Turk & Caprara, 2010) has also been implicated in mitochondrial genome stability (Guo, Chen et al., 2011, Shu, Vijayakumar et al., 2004, Tuteja, Tarique et al., 2014). Similar link between ribosome biogenesis with DNA replication has been found in both bacteria and yeast. In bacteria, Obg GTPases which are known to control large ribosomal subunit biogenesis (Datta, Skidmore et al., 2004, Jiang, Datta et al., 2006) links essential process required for cell cycle progression such as DNA replication by a yet unknown mechanism (Dutkiewicz, Slominska et al., 2002, Foti, Schienda et al., 2005, Kint, Verstraeten et al., 2014, Sikora, Zielke et al., 2006). Nog1 the nucleolar member of the Obg family of GTPases in yeast which is required for 60S ribosomal subunit biogenesis (Kallstrom, Hedges et al., 2003, Saveanu, Namane et al., 2003) is also associated with origin recognition complex in yeast (Berthon, Fujikane et al., 2009, Du & Stillman, 2002). Interestingly, we found Irc3 to be predominantly associated with the mitochondrial ribosomes in cells that are in logarithmic growth phase. However, in cells that have entered stationary phase, only a small fraction of Irc3 is associated with the mitochondrial ribosomes (Fig. 3). It would be interesting to speculate that Irc3 during rapid growth is involved in regulation of mitochondrial translation and switches to mtDNA maintenance at other times thus establishing a link between these two processes.

Mitochondrial functioning vis-à-vis ATP generation is altered depending on whether the cell is poised for rapid proliferation or in a quiescent/differentiated state. Proliferating cells, including pluripotent stem cells, use glycolysis even in presence of oxygen whereas differentiated cells use mitochondrial oxidative phosphorylation (Prigione, Ruiz-Perez et al., 2015, Vander Heiden, Cantley et al., 2009, Xu, Duan et al., 2013)*. Saccharomyces cerevisiae* shows a similar plasticity in response to metabolic cues i.e., when grown aerobically in glucose, yeast cells derive ATP from fermentation while producing ethanol as a by-product. Under these conditions the metabolism in yeast cells is akin to proliferating mammalian cells. Upon exhaustion of glucose yeast cells switch to oxidative phosphorylation for ATP generation from the ethanol produced during fermentation to sustain itself, akin to differentiated cells (Diaz-Ruiz, Rigoulet et al., 2011, Diaz-Ruiz, Uribe-Carvajal et al., 2009). We have shown that *IRC3* regulates mitochondrial translation distinctively in yeast mitochondria in response to metabolic cues. The closest orthologue for *IRC3* in humans based on BLAST is an uncharacterized putative RNA helicase (BAG50877) that shares 24% identity and 39% similarity within the helicase core only and predicted to localize to mitochondria with 84% probability (Claros & Vincens, 1996). Thus, although a clear orthologue for *IRC3* is not present in mammalian cells however, one could speculate that a similar mechanism exists in mammalian mitochondria to control synthesis of OXPHOS protein complex whereby energy demands and signals governing cellular state are integrated.

## MATERIALS AND METHOD

### Yeast strain, primers and media

Yeast strains used are listed in Table S1. Sequence of primers used are listed in Table S2. Complete media used were YEP (1% yeast extract and 2% peptone) containing 2% glucose (YPD), 2% galactose (YPGal) or 3% glycerol (YPG) as a carbon source. Synthetic minimal media (0.67% yeast nitrogen base without amino acids containing 2% glucose (SD), 2% galactose (SGal), 3% glycerol (SG), 0.1% 5-Fluoro-orotic acid, 2% glucose (5FOAD), 0.006% canavanine sulphate, 2% glucose (SCAN) were supplemented with appropriate amino acids when required, as described previously (Guthrie, 1991).

### Generation of plasmids and strains

To generate an episomally expressed *IRC3*, *IRC3* having 634bp’s upstream of start codon and 1kb downstream of stop codon in addition to ORF was PCR amplified using *YDR332w*up and *YDR332w*dn primer pair (Table S2) and cloned into pRS426 and pRS315 at the *Not*I site to generate *pRS426:IRC3* (pJk20) and *pRS315:IRC3* (pJk22). In order to generate a construct expressing *IRC3* c-terminally tagged Myc under its endogenous promoter pRS315::*IRC3-Myc,* Myc-KanMX6 was amplified from the pFA6a-13Myc-kanMX6 plasmid (Longtine, McKenzie et al., 1998) by using P3 and P4 primer pair (Table S2) and cloned into a pGEM-T easy vector at the TA cloning site to generate pJK55 (KDB784). *MycKanMX6* from pJK55 was subcloned into pJK34 (*IRC3::*3HA) as a *Bam*HI and *Sal*I fragment to generate pJK52 (KDB698).

Cell deleted for *IRC3* were generated in a diploid strain (KDY335) created by crossing CRY1 *(MATα; ura3-52; trp1Δ2; leu2-3_112; his 3-11; ade2-1; can 1-100)* and BY4742 *(Matα his3Δ1, leu2Δ0, lys2Δ0, ura3Δ0*). Disruption of *IRC3* was carried out by using a PCR-based gene replacement approach, where *HIS5* gene was amplified from pFAa-His3MX6 (Longtine et al., 1998)with upstream and downstream region of *IRC3* by using polymerase chain reaction using the oligonucleotide F1 and R1 (Table S2). The linear PCR product was transformed into diploid strain (KDY335) to create heterozygous diploid strain (KDY394). A plasmid borne *IRC3* (pJK20) was transformed into heterozygous diploid strain (KDY394), sporulated and haploid spores were selected on SCAN to generate a strain with chromosomal copy of *IRC3* deleted and Irc3 expressed from an episomal copy with intact mitochondrial DNA (KDY 495). In order to eliminate the chance of creating mutation at a second site in the genome during generation of *Δirc3::HIS5* and to possess mtDNA from an isogenic strain, KDY495 was back crossed six times with BY4741. The haploid strain generated upon back crossing was named as KDY1146. Replacement of wild type copy of *IRC3* with *HIS5* was similarly carried out in YC162, RG139, RG140 and W303ρ*I⁰* to generate KDY1575, KDY1529, KDY1531 and KDY1589 respectively.

In order to study translation initiation and elongation, *Δirc3* strain carrying ARG8^m^ reporter cassette was created by crossing KDY1146*ρ⁰*with XPM171a and XPM78a respectively. Diploid were sporulated and haploid spores were selected on SCAN, lacking arginine and histidine. To ensure that arginine gene from the nuclear genome in the haploids is disrupted, mitochondrial DNA was removed by growing cells in YPD supplemented with 25µg/ml Ethidium bromide (EtBr) and tested for growth on SD plates lacking arginine. The corresponding cell unable to grow on SD plates lacking arginine were selected from YPD plates and absence of *IRC3* gene from the nuclear background was confirmed by PCR using YDR332wup and YDR332wdn/R1 primer (Table S2) to generate KDY1447 and KDY1540.

### Presence of mitochondrial DNA

Presence of mitochondrial DNA in *Δirc3* expressing either *IRC3*(KDY1355) or empty vector (KDY 1366) were measured by crossing cells with *IRC3 ρ*⁰ testers followed by selection of diploid cells. The percentage of diploid cells that were able to utilize both glycerol and glucose was indicative of presence of mtDNA in the haploid strain.

To measure mitochondrial DNA copy number with respect to nuclear DNA in Δ*irc3* expressing either *IRC3* (KDY1355) or empty vector (KDY 1366) total yeast DNA was isolated and quantitative polymerase chain reaction (qPCR) was performed using primers COX2L and COX2R for Cox2 amplification and ActinF.P and ActinR.P for Actin amplification (Table S2). Ratio of signal for Cox2 to Actin was used as a measure of mitochondrial DNA to nuclear DNA respectively.

### Generation of temperature-sensitive (ts) allele of *IRC3*

Temperature sensitive *irc3* alleles were created using error-prone PCR as described previously (Stark, 1998). Coding region of was amplified by using tsmutantF.P and *YDR332w*dn primer pair (Table S2). PCR reactions were carried out in the presence of 5:1 ratio of (dTTP + dCTP):(dATP + dGTP) or (dATP + dGTP): (dTTP + dCTP). The concentration of MnCl_2_ was varied from 0.3 to 0.5 mM. The resulting mutated PCR fragment was cloned by gap repair into pJK22. Linearized pJK22 (digested with *Nsi*I) and mutagenized PCR product was co-transformed into KDY494. Transformed cells were selected on SD plates lacking leucine and uracil and patched on 5FOAD to negatively select for *URA3* helper plasmid expressing wild type allele of *IRC3*. Cells were then patched on glucose plates and incubated at 23⁰C and 37⁰C for 2 days. Cells were patched onto glycerol plates and scored for colony formation at permissive temperature (23⁰C) but not at non-permissive temperature (37⁰C). Plasmids DNA was isolated from Δ*irc3* cells harboring putative *ts* alleles and amplified in *E. coli* and sequenced. Two confirmed temperature sensitive allele of *IRC3* [pJK90 (KDB 1019) and pJK93 (KDB 1061)] were shuttled into KDY1146 and confirmed for growth at 23⁰C and not 37⁰C on YPG.

### Isolation of mitochondria and separation of mitochondrial ribosomes

Mitochondria were isolated from cells grown in YPGal, YPD or YPG at the indicated temperature at either OD_600_ of 1 or 3. Mitochondria was isolated by a previously described method (Glick & Pon, 1995) with slight modification described in (Datta et al., 2005). Mitochondrial ribosomes were separated by sucrose density gradient as described previously(Fearon & Mason, 1992). Mitochondria were resuspended in buffer D (10mM Tris-Cl, pH 7.4, 10mM magnesium acetate, 100mM NH_4_Cl, 7mM β-mercaptoethanol, and 1mM PMSF) and incubated on ice for 2hrs to generate mitoplasts. Mitoplasts were harvested by centrifugation at 12,000×g for 10 mins and resuspended in buffer D containing 2% NP 40, and further incubated on ice for 30 mins to allow swelling. Mitochondrial lysis was achieved by 20 strokes of tight-fitting pestle dounce homogeniser. The mitochondrial lysate was clarified by centrifugation at 40,000×g for 25 mins. Equivalent number of clarified lysates was loaded on 10-30% sucrose gradient containing 10mM Tris-Cl, pH7.4, 10mM magnesium acetate, 7mM β-mercaptoethanol and 400mM NH4Cl. Ribosomal particles were centrifuged at 135,000×g for 4hrs in a Beckman SW41Ti rotor. Equivalent fractions were collected and RNA contents were measured by UV absorbance at 254nm using an ISCO continuous flow cuvette. Protein samples in each fraction were precipitated using 15% TCA, separated by SDS-PAGE and subject to immunoblot analysis.

### Analysis of mitochondrial transcripts

Cells were grown in either YPD or YPGal media or shifted to YPG media. Cells were collected by centrifugation at 2,660×g for 5mins, washed with water and resuspended in SB3 buffer (50mM Tris-Cl pH8, 10mM MgCl_2_ 3mM DTT, 1M sorbitol). Spheroplast were generated by treating with zymolase (2.5mg/gm cell pellet) for 2 hrs at 37⁰C. Spheroplasts were collected by centrifugation at 4000xg for 5 mins and further lysed by 20 strokes of Dounce homogenizer. Cell lysate was centrifuged at 1,500xg for 5 mins at 4⁰C to remove cell debris. Supernatant was collected and centrifuged at 17,550xg for 15 mins at 4⁰C to pellet the mitochondria. Mitochondria was lysed by addition of trizol, followed by chloroform. The aqueous layer containing mitochondrial nucleic acid was separated and precipitated by addition of 3M sodium acetate and isopropanol. Pellet was washed twice with 70% ethanol to remove excess salts and resuspended in water. Isolated RNA was further purified by adding one volume of 2M potassium acetate pH 4.8, incubated on ice for 30 mins and centrifuged at 13,000 rpm, 4⁰C for 20 mins. Precipitated RNA was treated with DNase I (2000 Units/ml) for 30 mins at 37⁰C to remove contamination of nuclear or mtDNA, followed by precipitation with 3M sodium acetate and isopropanol. Precipitated RNA was washed twice with 70% ethanol and resuspended in water. cDNA using random primers was synthesized from 25ng/μl of RNA template. Synthesized cDNA was diluted 5 times prior to utilization in semi quantitative PCR reactions with 23 cycles extension for mitochondrial genes Cox1, Cox2, Cytb, Atp8, Atp9 and Atp6 using primers listed in Table S2. For clarity, agarose gel pictures shown in the figures were colour-inverted, resulting in dark bands on a white background.

### Analysis of mitochondrial protein synthesis

Cell were grown in SD or SGal or SG lacking methionine and cysteine at the indicated temperature to an O.D_600_ of 1. Newly synthesized mitochondrial proteins were labelled with 0.1mCi of [^35^S]methionine and cysteine [Easy Tag^tm^ Express^35^S protein labelling mix, Perkin Elmer NEG77200 (specific activity: 1175 Ci/mmol)] in the presence of cycloheximide for 30 mins as described in (Fox, Folley et al., 1991). Mitochondria was isolated from labelled cells as described previously (Fox et al., 1991). Mitochondrial proteins were quantified by Bradford assay and equivalent amounts of protein were separated on 17.5% SDS-PAGE. Labelled proteins were either transferred to nitrocellulose membrane and analysed by phosphoimager FLA9000 or gels were dried and exposed of X-ray film to visualize labelled proteins. Coomassie stained gels were used as a part of normalization of radiolabelled mitochondrial proteins.

### Immunoblot Analysis

Proteins were separated on 7.5% or 10% SDS-PAGE and subjected to immunoblot analysis. Following antibodies were used: anti-Myc (1:5000) (MYC.A7; Epitope Biotech Inc.), Mrp7 (1:200) (Fearon & Mason, 1988), Mrp13 (1:200) (Partaledis & Mason, 1988) and Arg8p (Steele et al., 1996), Mtg2 (Datta et al., 2005).

## Acknowledgements

We are extremely grateful to Prof. Janine R. Maddock for strains, plasmids and antibodies, Prof. Thomas D. Fox for antibodies to Arg8, Prof. Xochitl Pérez Martínez for strains YC162, RGV139, RGV 140 and XPM78a and Prof. Antoni Barrientos for strain XPM171a. We also thank Prof. Jagreet Kaur and Prof. Suman Dhar for critical reading of this manuscript. We also thank instrumentation facilities at CIF, UDSC and DST-FIST/UGC-SAP supported CIF, Genetics. This work was supported by grants from Department of Biotechnology (Grant numbers: BT/PR14740/BRB/10/875/2010, BT/PR15104/GBD/27315/2011), SERB (EMR/2015/000650) and CSIR (27(0324)/EMR-II) and R&D grant from Delhi University to K.D. J.K. acknowledges UGC, BSR, SERB and ICMR for junior research fellowship and senior research fellowships.

## Author Contributions

J.K. performed experiments and analysed data. K.D. and J.K. conceived the project, designed experiments, analysed the data and wrote the paper. K.D. was responsible for securing funding for this work.

## Conflict of interest

The authors declare that they have no conflict of interest.

## Supplementary Figures

**Fig. S1:**
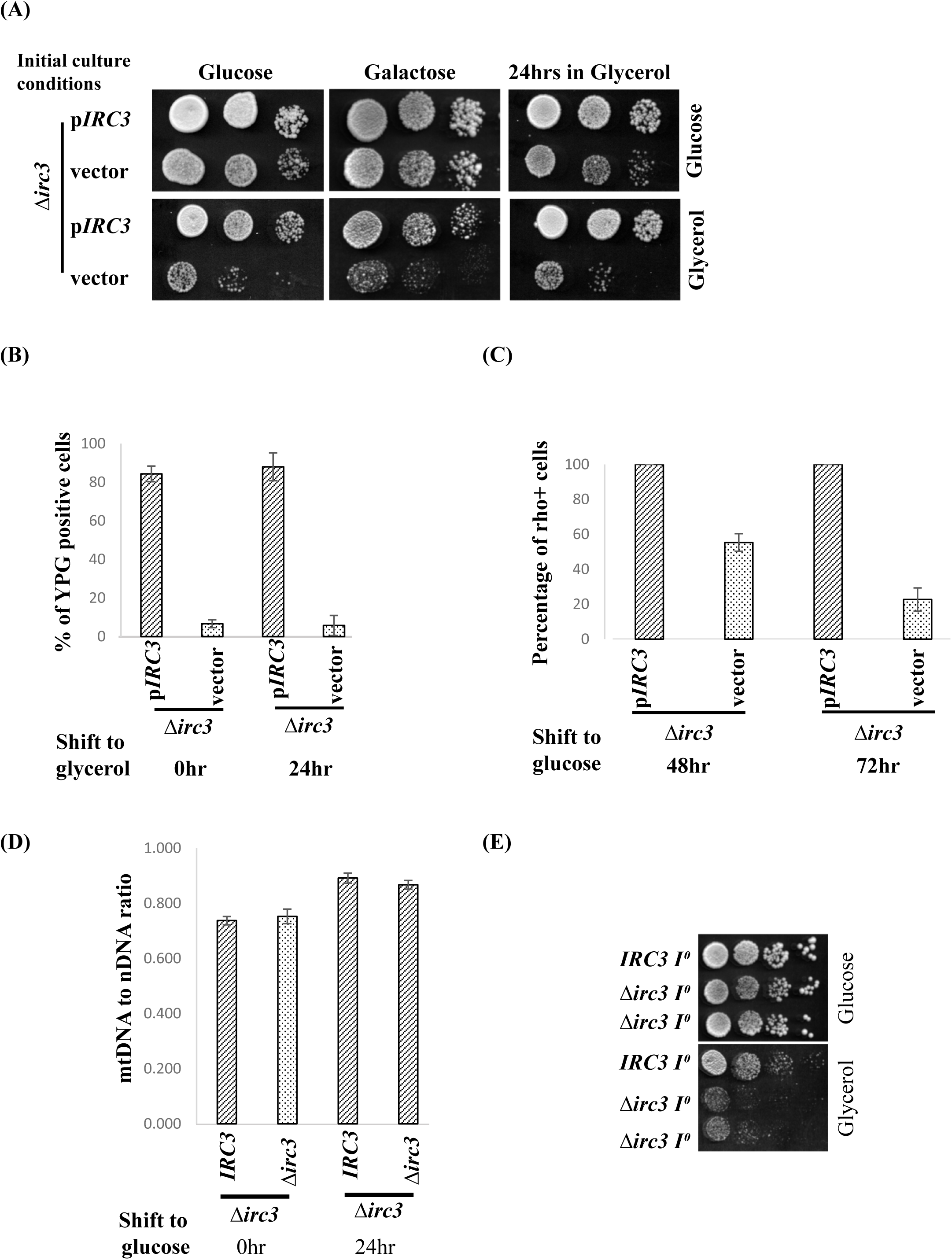
Loss of growth on glycerol precedes loss of mtDNA in Δ*irc3* cells due to aberrant gene expression. (A) Shown are tenfold serial dilution of Δ*irc3* cells expressing either wild type *IRC3* or vector cultured in glucose, galactose and 24hrs in glycerol followed by spotting on glucose and glycerol plates. (B) Percentage of Δ*irc3* cells expressing either wild type *IRC3* or empty vector that are able to utilize glycerol as the carbon source upon subculturing in glycerol media for indicated time period. (C) Percentage of mtDNA present in Δ*irc3* cells expressing either wild type *IRC3* or vector at the indicated time upon subculturing in glucose. The percentage of *ρ^+^* cells obtained from initial time points are indicated in (Fig. 1C). (D) Ratio of nuclear to mitochondrial DNA, was estimated Δ*irc3* cells expressing either wild type *IRC3* or vector at the indicated time after subculturing in glucose.(E) Shown are tenfold serial dilution of wild type and Δ*irc3* cells carrying intron less mitochondrial DNA (*I⁰*) on glucose and glycerol plate.

**Fig. S2:**
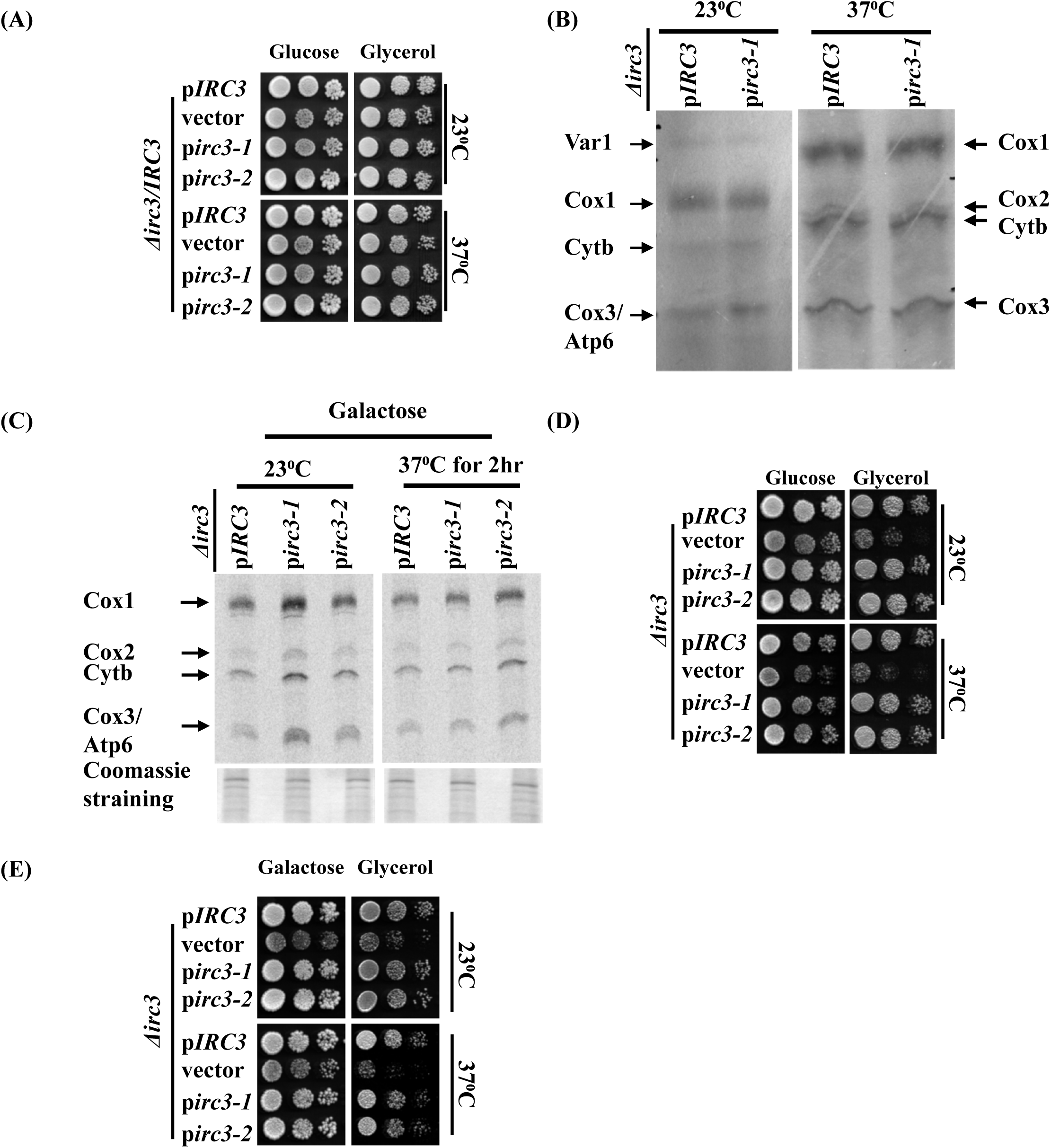
Depletion of functional *IRC3* in cells cultured in galactose/glycerol do not alter mitochondrial protein synthesis. (A) Shown are tenfold serial dilutions of diploids generated by crossing *IRC3ρ⁰* tester strain with Δ*irc3* cells expressing either wild type *IRC3,* vector *irc3-1* or *irc3-2* grown in glucose at 23⁰C and 37⁰C for 4 days. Presence of mitochondrial DNA was determined as a function of diploid cells ability to utilise glycerol as a sole carbon source. Mitochondrial protein synthesis was measured in the indicated strains cultured in (B) glycerol or (C) galactose either at 23⁰C or upon shift to 37⁰C for 2hrs by incorporation of [^35^S]methionine and cysteine in the presence of cycloheximide. Mitochondria from labelled cells were isolated and proteins were separated on 17.5% SDS-PAGE. Radiolabeled proteins were visualized by autoradiography and phosphoimaging. Below are Coomassie stained gel of radiolabeled mitochondrial extracts. (D) Shown are tenfold serial dilution of the indicated strains cultured in glycerol for 24hrs hours prior to spotting on glucose and glycerol plates and further incubated at 23⁰C or 37⁰C. (E) Shown are tenfold serial dilution of the indicated strains cultured in galactose were spotted on galactose and glycerol plates, followed by incubation at 23⁰C and 37⁰C for 2 days.

**Fig. S3:**
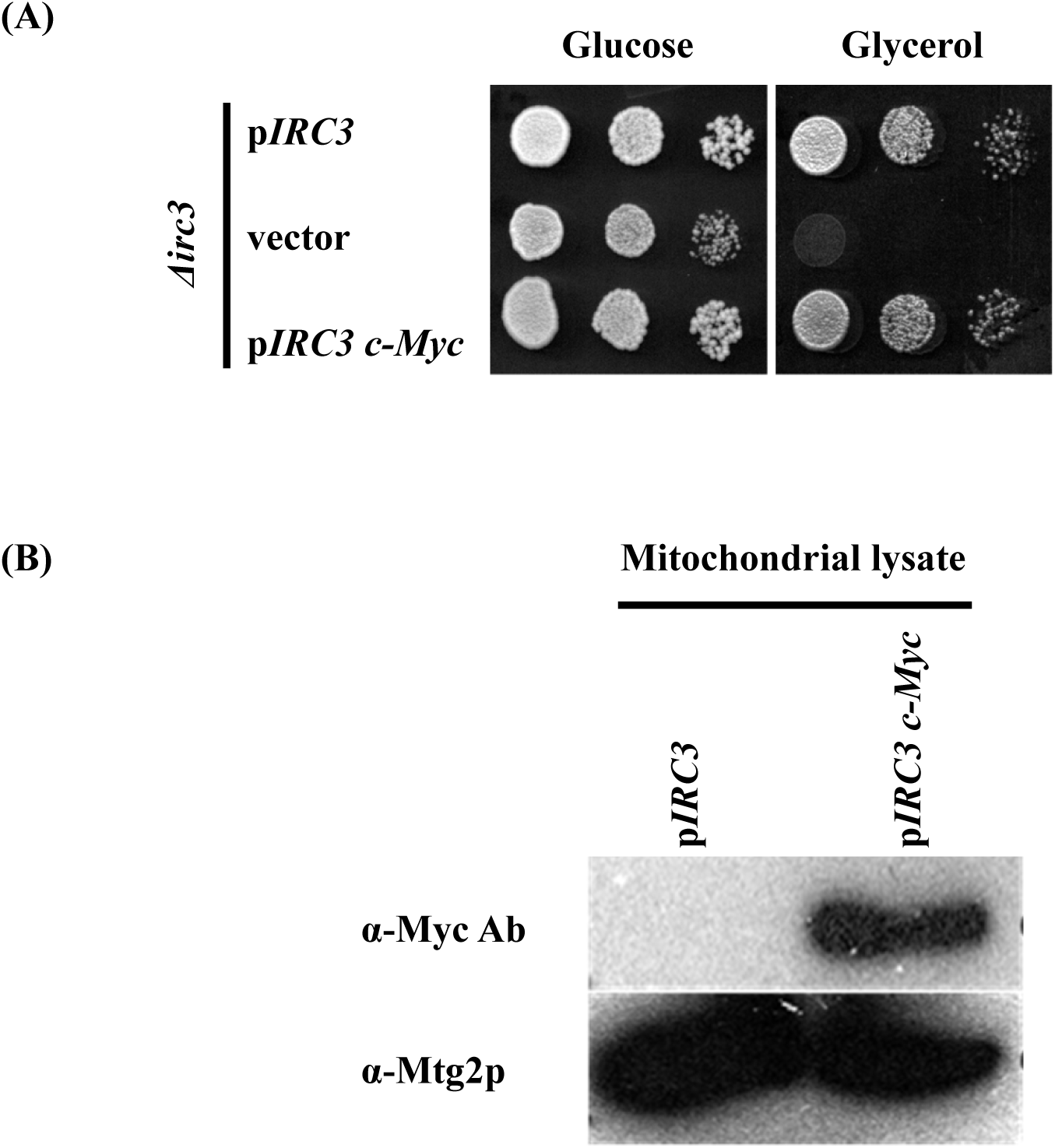
Irc3-Mycp encodes a functional protein. (A) Shown are tenfold serial dilution of Δ*irc3* cells expressing either wild type *IRC3,* vector or *IRC3c-Myc* spotted on glucose and glycerol plates. (B) Mitochondrial extract prepared from Δ*irc3* cells expressing either *IRC3* or *IRC3c-Myc* were subjected to immunoblot analysis using anti-Myc and anti-Mtg2 antibodies.

**Fig. S4:**
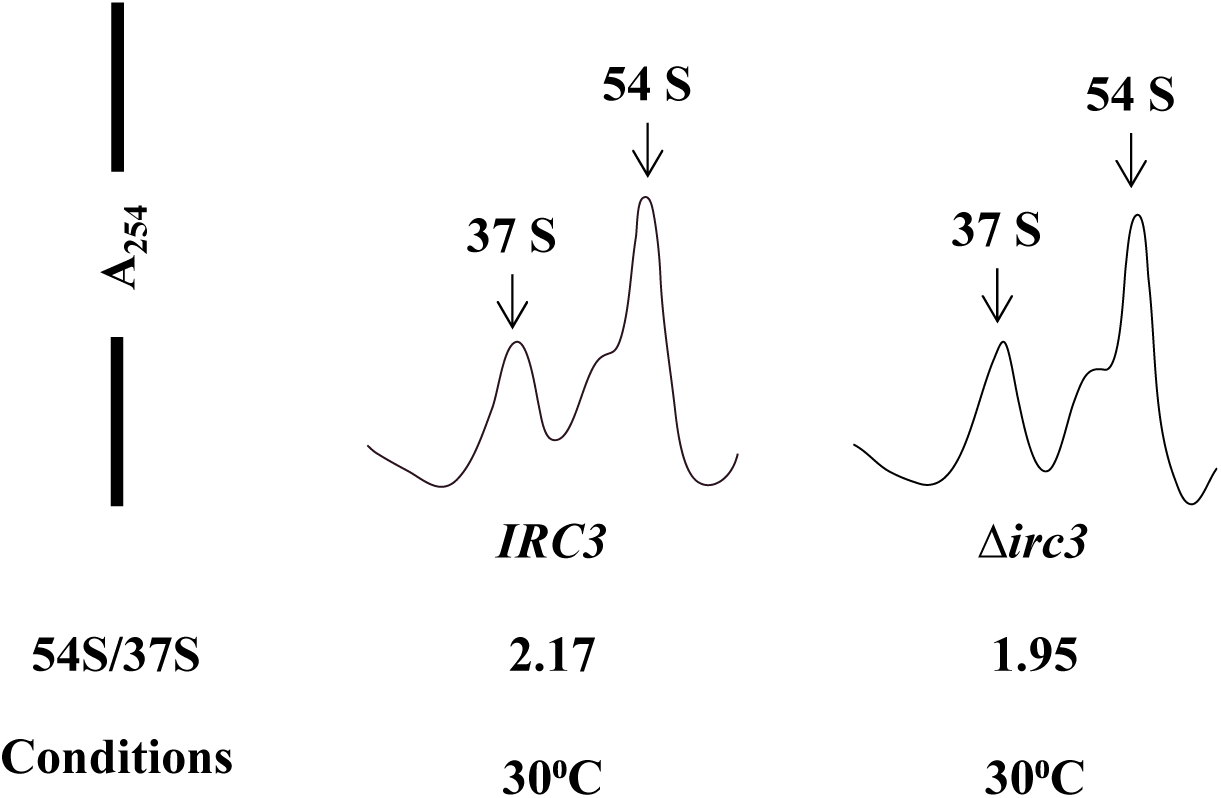
Mitochondrial ribosomal subunit is not altered in Δ*irc3* cells cultured in glycerol. Mitochondrial ribosomal subunits from Δ*irc3ρ^+^* cells expressing either wild type *IRC3* or vector, grown in glycerol medium for 12hrs, were fractionated on 10-30% sucrose gradient. RNA content was monitored at 254nm. The position of 37S and 54S peaks are labeled. The relative ratio of 54S/37S peaks are listed below each profile.

**Fig. S5:**
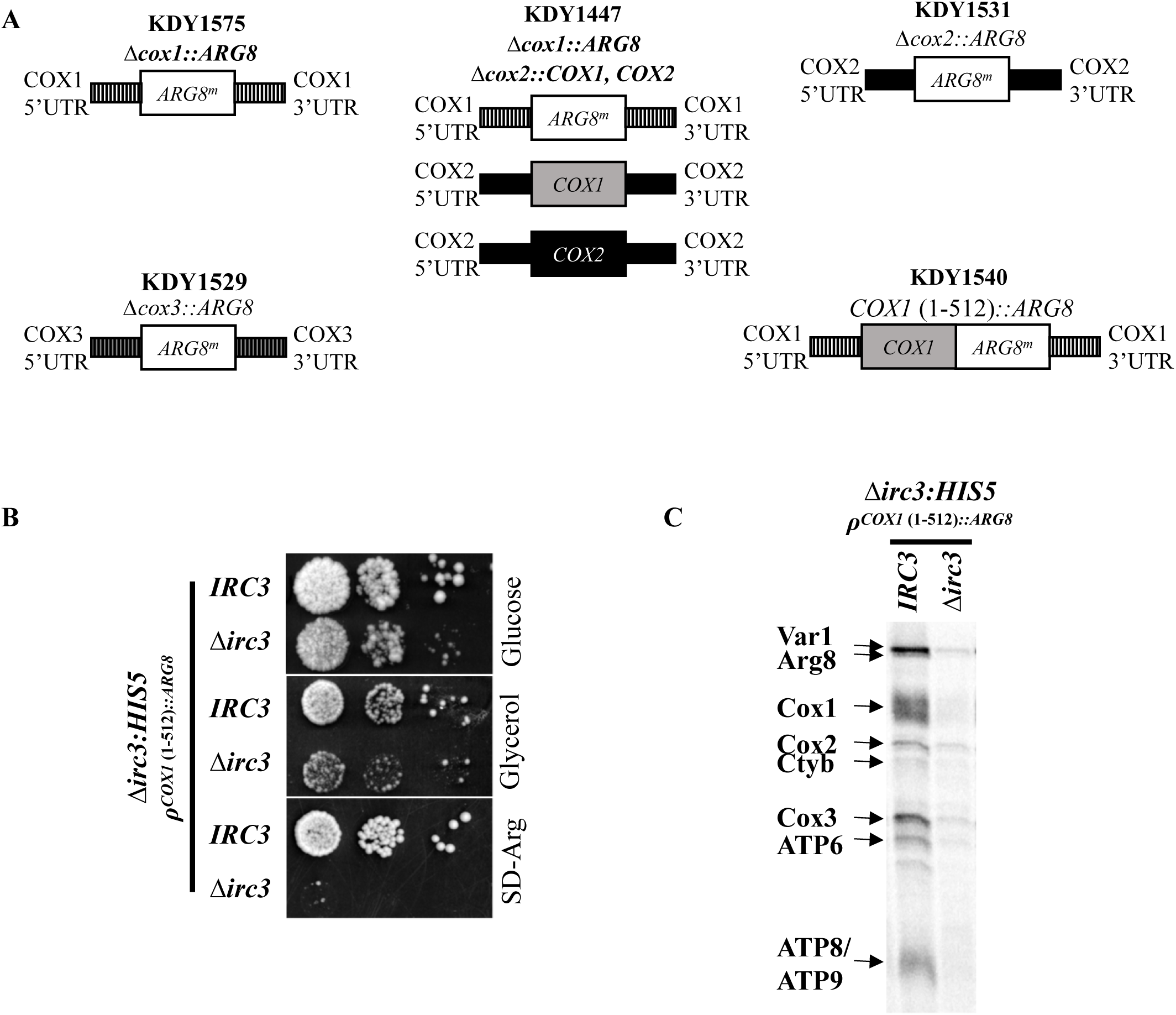
Irc3 is required for mitochondrial translation in engineered strains. (A) Schematic representation of mitochondrial genotype of strain (KDY1575, KDY1529, KDY1531, KDY1447 and KDY1540) used to study translation initiation or elongation that are derived from YC162, RGV140, RGV139, XPM171a and XPM78a respectively. (B) Shown are tenfold serial dilution of Δ*irc3* cells in XPM78a (KDY1540) harboring an episomal *IRC3* allele or vector were spotted on YPD, SD-Arg and YPG (C) Mitochondrial protein synthesis was measured inΔ*irc3* cells in XPM78a harboring an episomal *IRC3* allele or vector at 23⁰C by incorporation of [^35^S]methionine and cysteine in the presence of cycloheximide. Mitochondria from labelled cells were isolated and proteins were separated on 17.5% SDS-PAGE. Radiolabeled proteins were visualized by phosphoimaging.

**Fig. S6:**
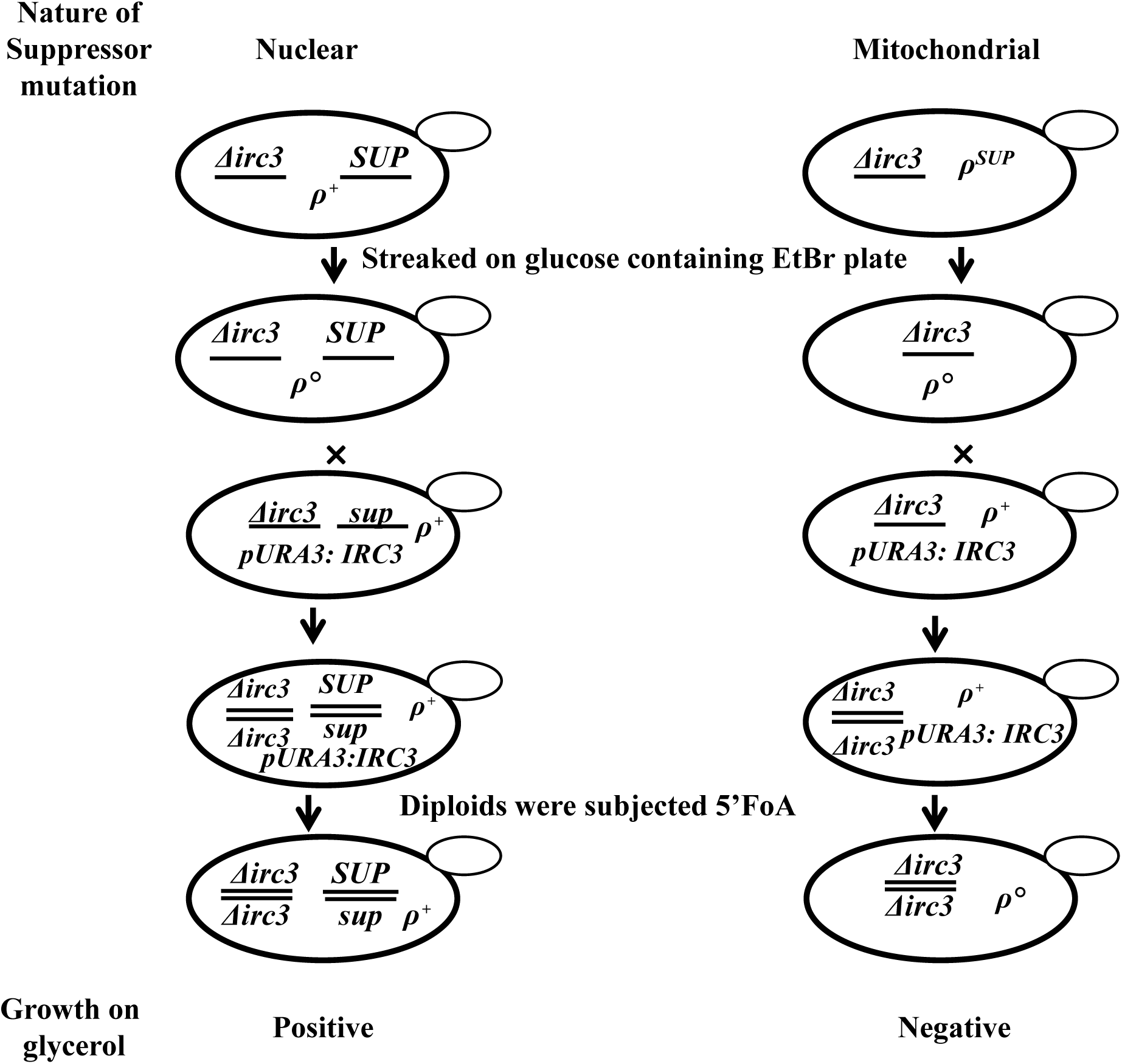
Schematic for differentiating whether suppressor mutation (sup) has mitochondrial or nuclear inheritance. Suppressors were plated on glucose plate containing ethidium bromide to generate *ρ⁰*derivatives, followed by mating with KDY495 (Δ*irc3/pRS426::IRC3 ρ^+^*). Diploids were plated on 5’FOAD to negatively select *pRS426::IRC3* and spotted on glucose and glycerol plates. The ability of diploids to utilise glycerol was indicative of the type of mutation.

## Supplementary Tables

**Table S1:**
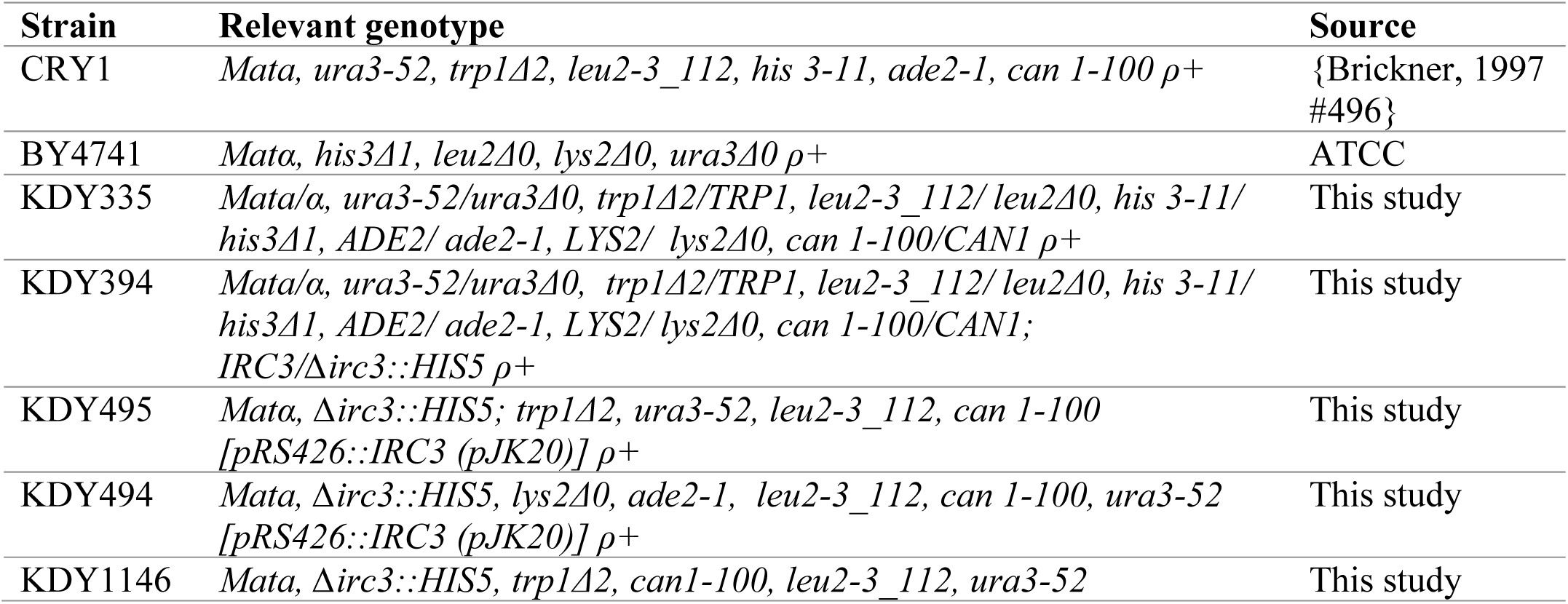
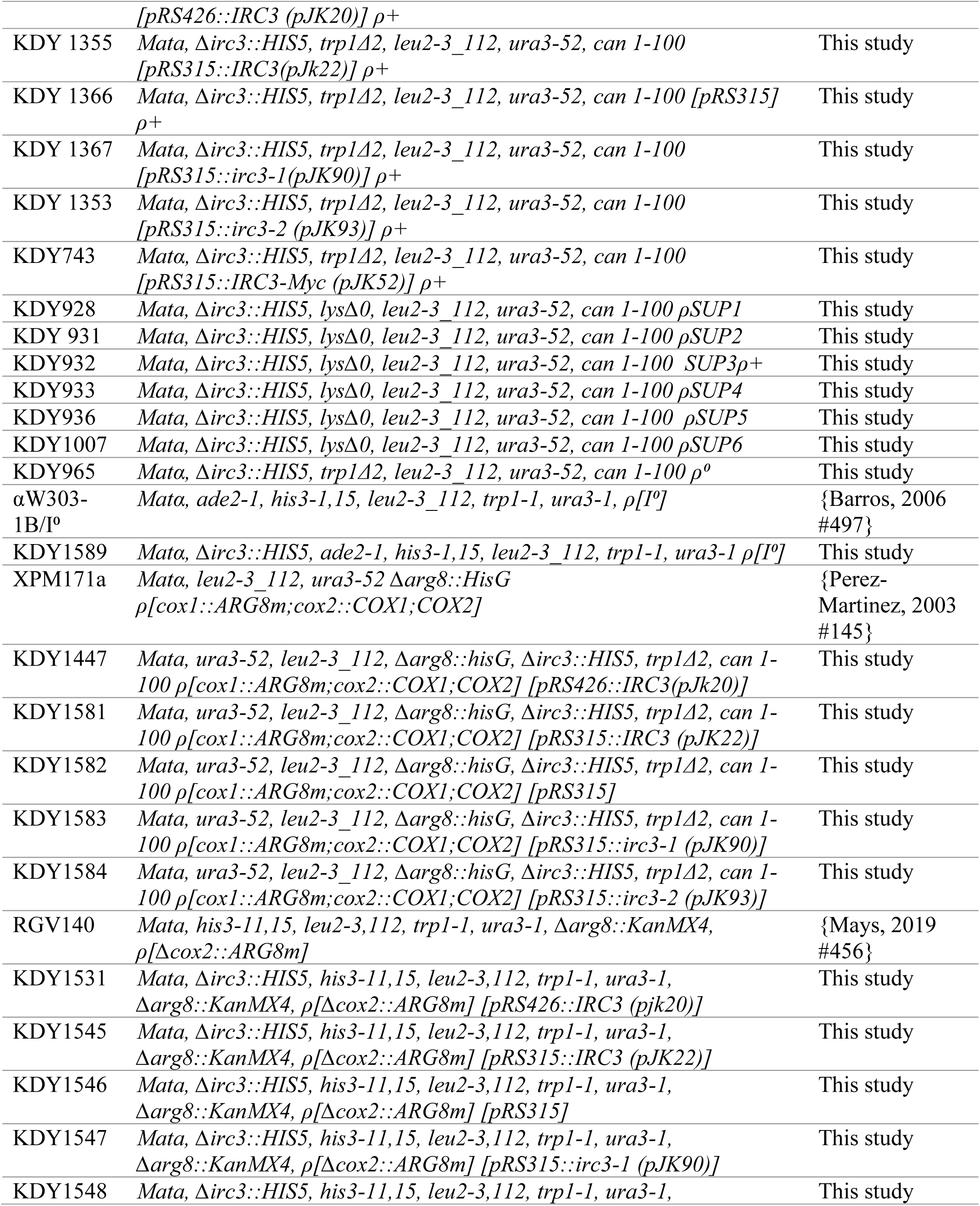
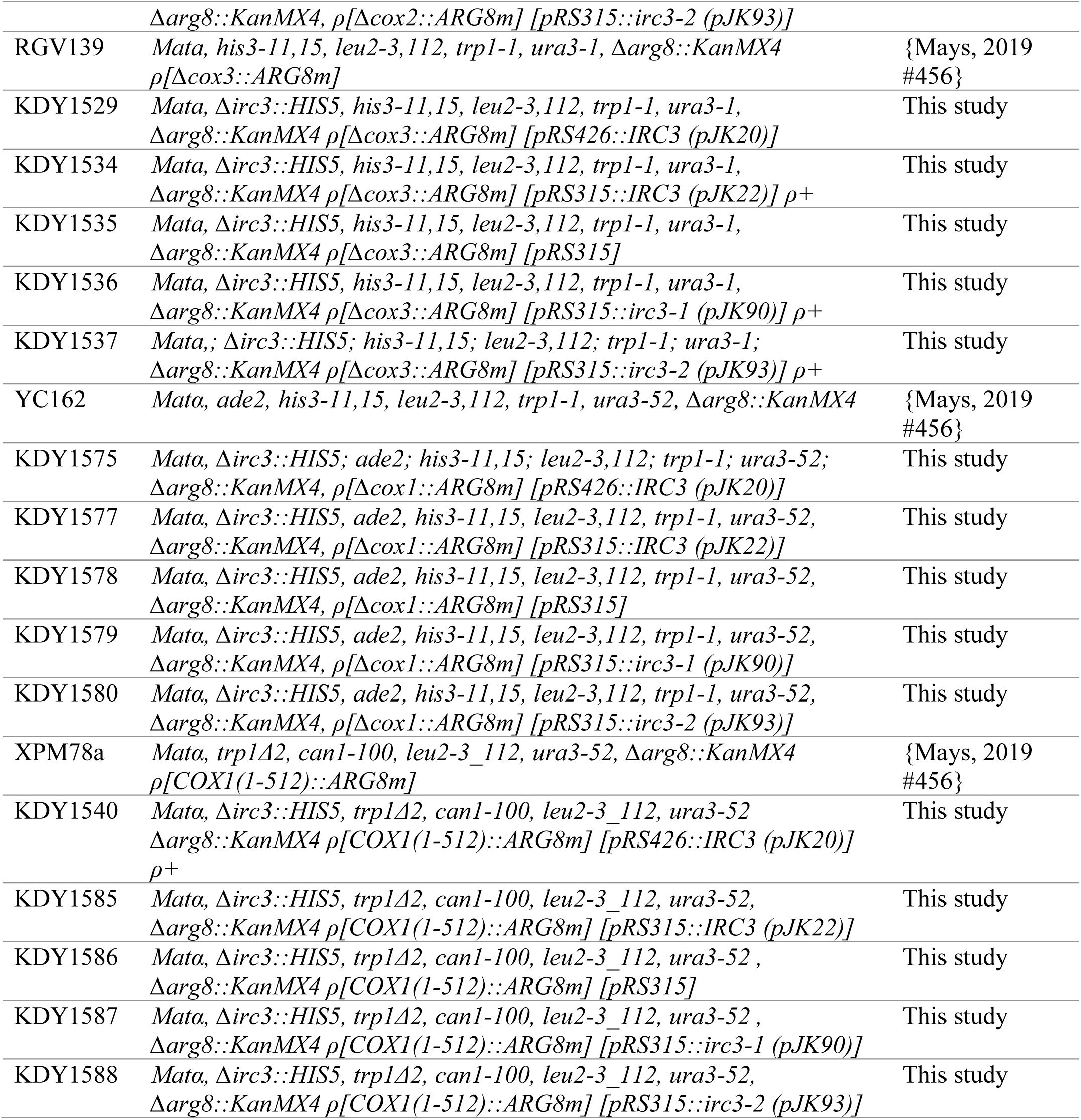
Yeast strain used in this study

**Table S2:**
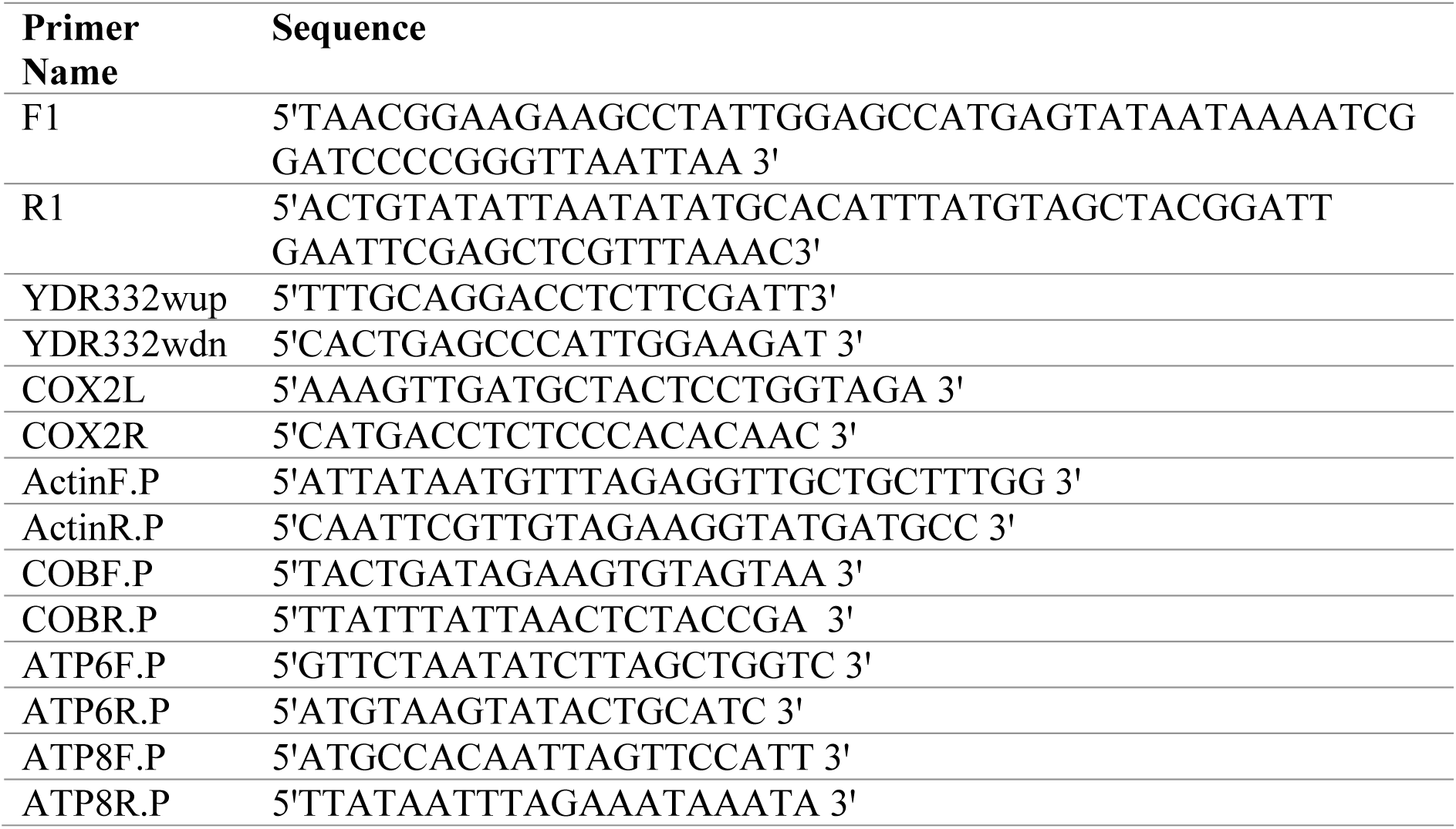
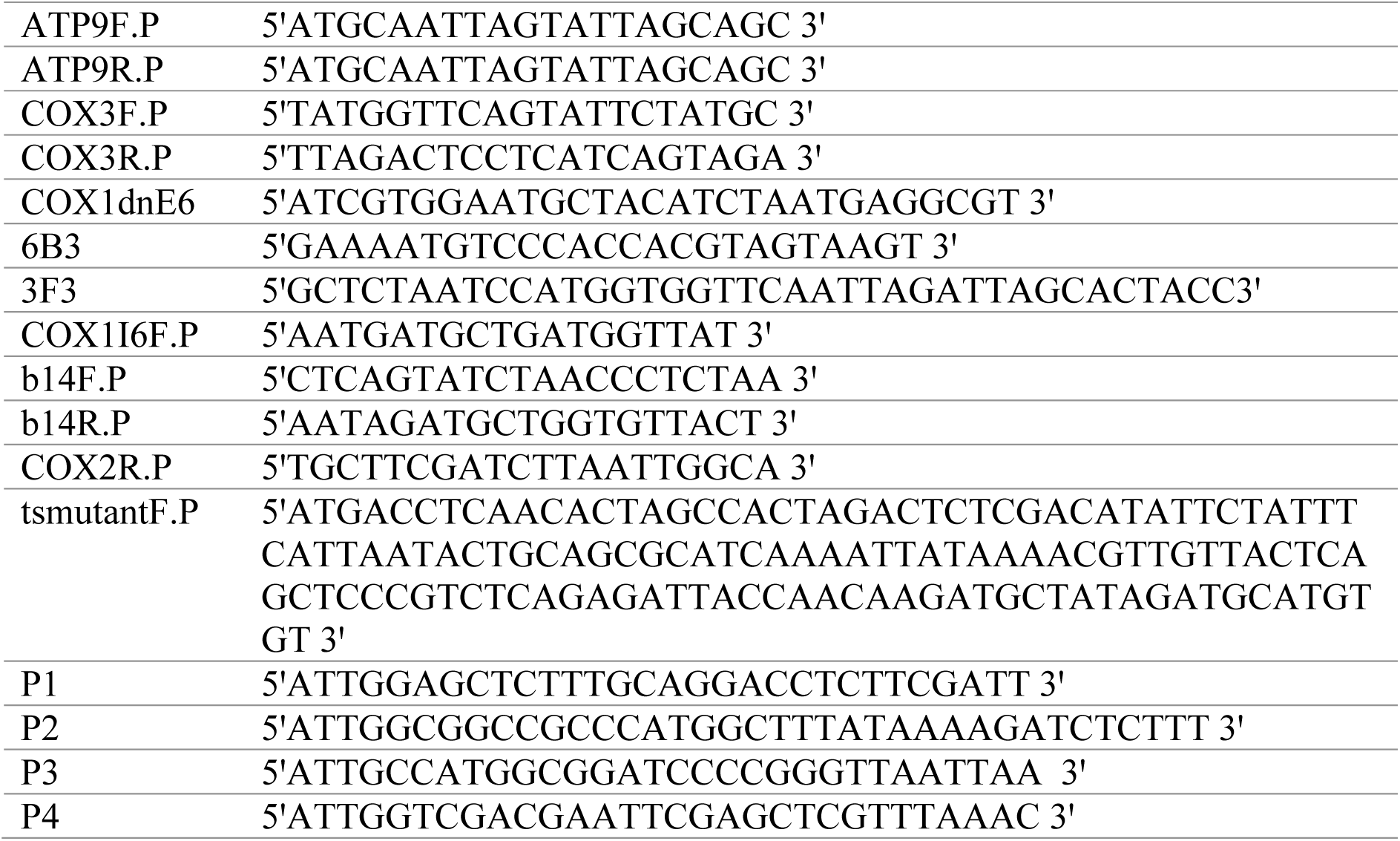
List of primers used in this study

**Table S3.**
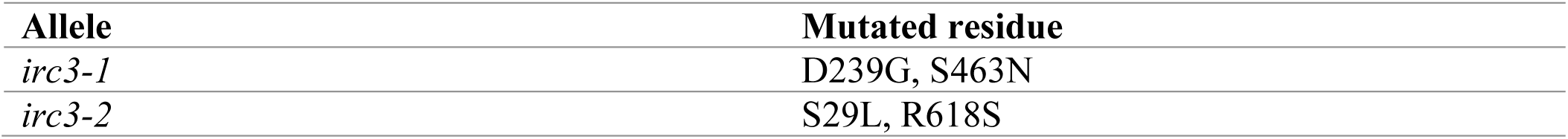
The mutated residues in temperature sensitive alleles of *irc3*

| Allele | Mutated residue |
| --- | --- |
| *irc3-1* | D239G, S463N |
| *irc3-2* | S29L, R618S |

